# More than just front or back: Parietal-striatal-thalamic circuits predict consciousness level

**DOI:** 10.1101/2020.04.07.030429

**Authors:** Mohsen Afrasiabi, Michelle J. Redinbaugh, Jessica M. Phillips, Niranjan A. Kambi, Sounak Mohanta, Aeyal Raz, Andrew M. Haun, Yuri B. Saalmann

**Affiliations:** Department of Psychology, University of Wisconsin-Madison, Madison, WI, 53705, USA; Department of Anesthesiology, Rambam Health Care Campus, Haifa, 3109601, Israel and The Ruth and Bruce Rappaport Faculty of Medicine, Technion – Israel Institute of Technology, Haifa 3200003, Israel; Department of Psychiatry, University of Wisconsin-Madison, Madison, WI, 53705, USA; Wisconsin National Primate Research Center, Madison, WI, 53705, USA

**Author notes:** Correspondence to: Yuri Saalmann, 1202 W Johnson St, Madison, WI 53706, Mohsen Afrasiabi, & Michelle Redinbaugh. These authors contributed equally (Mohsen Afrasiabi, Michelle J. Redinbaugh).

## Abstract

Major theories of consciousness disagree on the key neural substrates. In Global Neuronal Workspace Theory and Higher-order Theories, consciousness depends on frontal cortex, whereas Integrated Information Theory and Recurrent Processing Theory highlight posterior contributions. Most theories omit subcortical influences. To test these theories, we performed simultaneous frontal, parietal, striatal and thalamic recordings from awake, sleeping and anesthetized macaques, further manipulating consciousness with deep-brain thalamic stimulation. Information theoretic measures and machine learning approaches suggested parietal cortex, striatum and thalamus contribute more to consciousness level than frontal cortex. While these findings provide greater support for Integrated Information Theory than the others, the theory does not incorporate subcortical structures such as the striatum. We therefore propose that thalamo-striatal circuits have a cause-effect structure to generate integrated information.

**One Sentence Summary:** Parietal, but not frontal, circuits incorporating striatum and thalamus predict consciousness.

## Main Text

Consciousness is the capacity to experience one’s environment and internal state. Major theories disagree about the neural correlates of consciousness (NCC), notably the causal contributions of frontal relative to more posterior cortex, thus framing debates as “front versus back”. Both Global Neuronal Workspace Theory (GNW; Fig. S1B) and Higher-order theories (HOT; Fig. S1C) emphasize the role of frontal cortex. In GNW, consciousness depends on an “ignition” process in which information becomes globally available through top-down signals from frontal cortex (*1, 2*). In HOT, consciousness depends on frontal cortex supporting a higher-order thought about a sensory experience (*3, 4*). In contrast, Integrated Information Theory (IIT; Fig. S1D) and Recurrent Processing Theory (RPT; Fig S1E) emphasize the role of posterior cortex. For IIT, consciousness depends on cause-effect structures generating integrated information (*5*); proponents have claimed a posterior hot zone, including parietal cortex, to be the most critical (*6*). For RPT, consciousness depends on recurrent activity between earlier and later stages in sensory pathways (*7*). Some functional MRI and electrophysiology studies (*8, 9*) have suggested contributions of frontal cortex to conscious awareness of sensory stimuli, though critics claim these frontal responses may reflect report or executive functions (*10, 11*). Lesion studies have suggested consciousness persists despite large-scale frontal damage (*10*), though critics claim the lesions are incomplete (*12*). These concerns highlight the need for evidence based on no-report paradigms in healthy subjects (*11*).

Although important, these debates are unnecessarily cortico-centric. The higher-order thalamus, which robustly interconnects with the entire cortex, supports information transmission and recurrent activity between cortical neurons (*13-15*). Though the thalamus is under inhibitory control from the basal ganglia, some have used the striatum as an example brain area that does not directly contribute to consciousness, because of the common view of unidirectional information flow from striatum to thalamus (*10, 16*). However, as direct projections from intralaminar and other thalamic nuclei to the striatum allow bidirectional flow (*17, 18*), and the striatum widely samples the cortex, others have proposed a striatal role in consciousness (*19*).

To probe contributions of thalamus, striatum, frontal (F) and parietal (P) cortex to consciousness, we used linear electrode arrays to simultaneously measure local field potentials (LFPs) in superficial, middle and deep layers of the right frontal eye field (F_s_, F_m_ and F_d_, respectively), lateral intraparietal area (P_s_, P_m_ and P_d_), central lateral thalamus (T) and caudate nucleus (C) of macaques (constituting 8 parts of our cortico-striatal-thalamic system; Fig. S1A). Recordings occurred in three states – general anesthesia (propofol, isoflurane), light non-rapid eye movement (NREM) sleep, and resting (no report) wakefulness in a dark, quiet room – and two stimulation conditions – effective, in which we stimulated T to rouse monkeys from stable anesthesia, and control, which did not change consciousness. We used spectral power and three information theoretic measures – entropy (*H*), mutual information (*I*), and integrated information (Φ*; Fig. S2) – to decode the state of consciousness (*20*). We measured the differential weights across classifier features to probe their contributions to consciousness.

All measures showed state-dependent variation. LFP power varied by state, area and frequency: there was greater delta (1-4 Hz) power for anesthesia (all areas, t(263) ≥ 4.54, p ≤ 8.62×10^−6^) and light NREM sleep (frontal cortex, t(789) = 2.69, p = .029) relative to wakefulness (Fig. 1A). Φ* (Fig. 1B; see Fig. S3 for controls), *I*, and *-* (Fig S4, A and D) were higher in wakefulness relative to sleep and anesthesia (t(354) ≥ 2.54, p ≤ .011). To test how well each measure related to consciousness, we built classifiers to decode conscious state (wake, sleep, anesthesia) from LFP power (48 features: 8 system parts x 6 frequencies), Φ*, *I* or *H* (9 features: 8 n-1 subsystems plus 1 full system of n=8 parts). All classifiers decoded better than chance (33%; Fig. 1C; t(20) ≥ 5.42, p ≤ 3.14×10^−5^). Of these classifiers, Φ* (92.8%) was significantly more accurate (t(80) ≥ 4.73, p ≤ 2.06×10^−5^), followed by the LFP power classifier (t(80) ≥ 5.75, p ≤ 5.37×10^−7^). The Φ* classifier performed better for wake (F-Score_W_ = 0.95), sleep (F-Score_S_ = 0.93) and anesthesia (F-Score_A_ = 0.97) than the LFP classifier (Fig. 1, D and E; F-Score_W_ = 0.79, F-Score_S_ = 0.78, F-Score_A_ = 0.87). Φ* yielded greater accuracy for decoding conscious state, despite the classifier having fewer features.

**Fig. 1.**
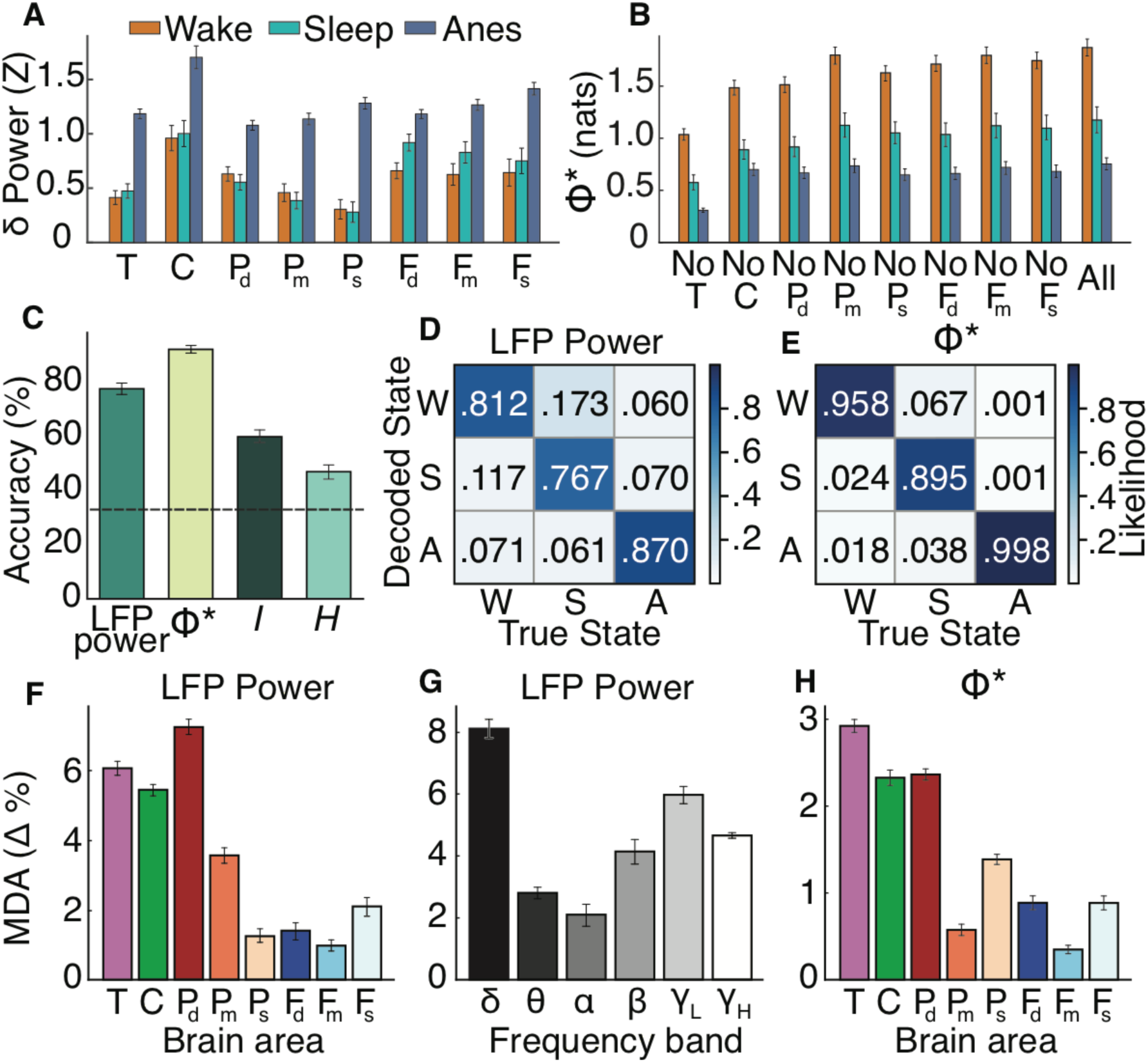
Parietal deep layers and subcortical areas contribute most to state decoding. (**A**) Normalized average delta power (Z score ± SE) for thalamus (T), caudate (C), and superficial (s), middle (m) and deep (d) layers of frontal (F) and parietal (P) cortex during wakefulness, sleep and anesthesia (Anes). (**B**) Population mean Φ* (± SE) during different states for the full system (All) or subsystems with one area removed. (**C**) Decoding accuracy (± SE) for classifiers using LFP power, Φ*, mutual information (*I*) or entropy (*H*). Dashed line shows chance. (**D**) Confusion matrix of LFP power classifier for wakefulness (W), sleep (S) and anesthesia (A); color scales with classification likelihood. **(E)** Confusion matrix of Φ* classifier. (**F-H**) Mean decrease in accuracy (MDA ± SE) for LFP power classifier after removing features for (**F**) brain areas or (**G**) frequency bands, or (**H**) Φ* classifier after removing subsystems containing specified brain area.

Because there was limited dependence between dissimilar features (Fig. S5, A and B; (*21*)), we computed the mean decrease in accuracy (MDA) after removing certain features as a measure of their importance. For the LFP power classifier, removing P_d_, T and C features caused larger MDAs than other areas (P_m_, P_s_, F_d_, F_m_, F_s_; t(160) ≥ 12.11, p ≤ 4.8×10^−15^; Fig. 1F; Table S1). In the frequency domain, removing delta and low-gamma (30-60 Hz) features caused larger MDAs than other bands (θ = 4-8 Hz, α = 8-15 Hz, β = 15-30 Hz, γ_H_ = 60-90 Hz; t(240) ≥ 5.99, p ≤ 2.36×10^−8^; Fig. 1G; Table S1). We performed similar analyses for the Φ* classifier (Fig. 1H), and MDAs were significantly greater after removing subsystems with P_d_, T or C over other parts (P_m_, P_s_, F_d_, F_m_, F_s_; |t(80)| ≥ 9.12, p ≤ 1.76×10^−12^; Table S1). These results show the importance of parietal and subcortical areas over frontal cortex.

We previously reported that 50 Hz thalamic stimulation could increase the consciousness level (measured via 10-point scale; (*21*)) of monkeys under stable anesthesia relative to control (ineffective) stimulation (*22*). To assess decodability of stimulation conditions, we calculated LFP power, *I, H* and Φ* prior to, during and after effective (consciousness level M = 4.37, SD = 1.52, n = 35) and control (M = 0.66, SD = 0.99, n = 128) stimulations. For all areas, LFP delta power tended to increase (Fig. 2, A and B) during thalamic stimulation (t(332) ≥ 3.58, p ≤ 4.01×10^−4^), and did not interact with effectiveness (|t(332)| ≤ .96, p ≥ .34). Results were similar for other frequency bands. However, Φ* showed clear increases selective to effective stimulations (Fig. 2, C and D); the interaction was significant (t(326) = 4.03, p = 2.06×10^−4^). There was no significant interaction (|t(326)| ≤ .80, p ≥ .35) for *H* (Fig. S4, B and C) or *I* (Fig. S4, E and F).

**Fig. 2.**
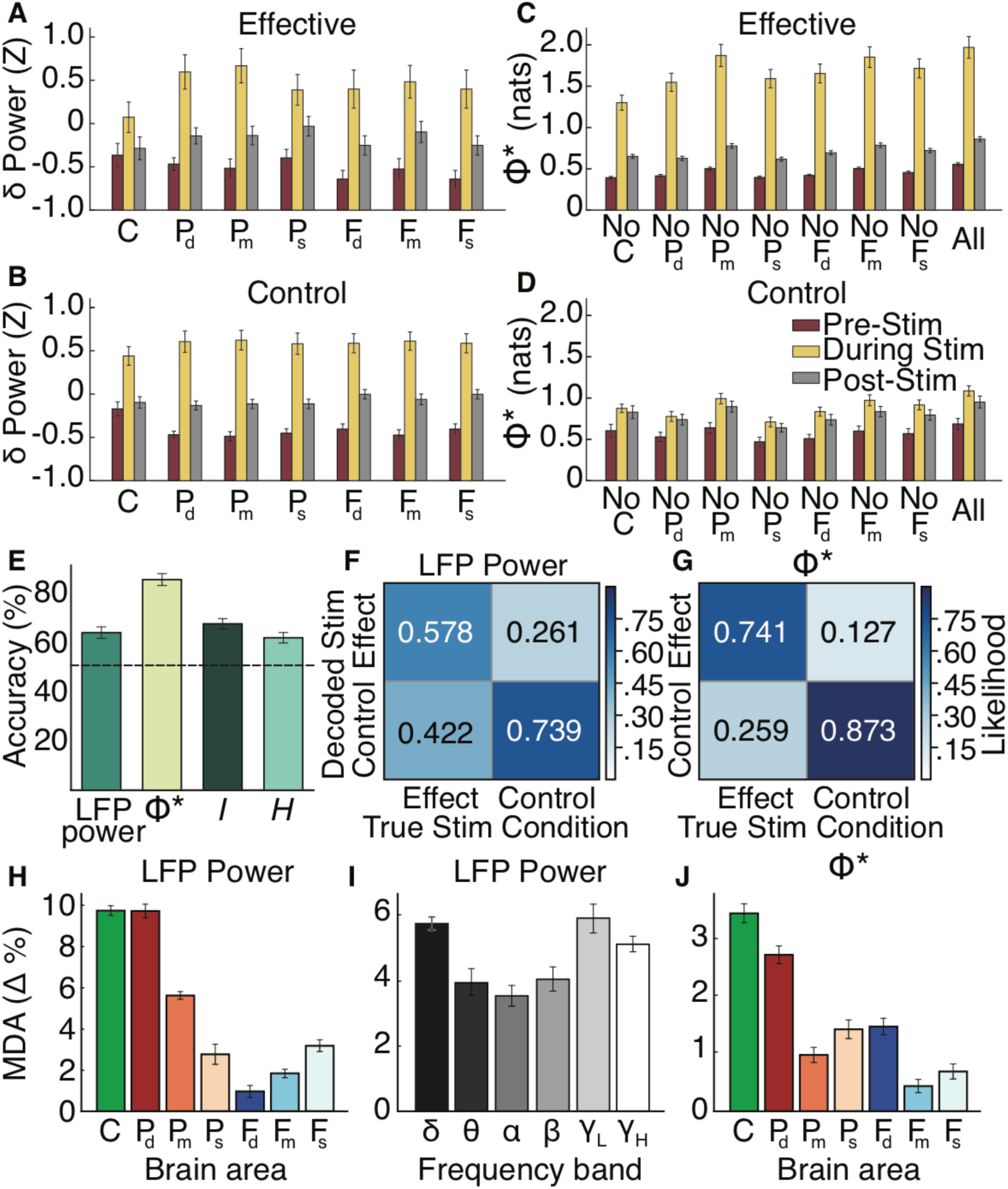
Parietal deep layers and subcortical areas contribute most to decoding stimulation-induced consciousness. (**A** and **B**) Normalized average delta power (Z score ± SE) for caudate (C) and superficial (s), middle (m) and deep (d) layers of frontal (F) and parietal (P) cortex prior to, during and after (**A**) effective or (**B**) control thalamic stimulations. (**C** and **D**) Population mean Φ* (± SE) for the full system (All) or subsystems with one area removed surrounding (**C**) effective or (**D**) control stimulations. (**E**) Decoding accuracy (± SE) for classifiers using LFP power, Φ*, mutual information (*I*) or entropy (*H*). Dashed line shows chance. (**F**) Confusion matrix of LFP power classifier for effective (Effect) or control stimulations; color scales with classification likelihood. (**G**) Confusion matrix of Φ* classifier. (**H-J**) Mean decrease in accuracy (MDA ± SE) for LFP power classifier after removing features for (**H**) brain areas or (**I**) frequency bands, or (**J**) Φ* classifier after removing subsystems containing specified brain area.

To test how each measure related to consciousness driven by thalamic stimulation, we built classifiers to decode induced conscious state (effective, control) from LFP power (42 features: 7 system parts x 6 frequencies), Φ*, *I* and *H* (8 features: 7 n-1 subsystems plus 1 full system of n=7 parts); thalamus not recorded during stimulation. All classifiers decoded better than chance (50%; Fig. 2E; t(20) ≥ 5.32, p ≤ 3.95×10^−5^). Of these classifiers, Φ* (84.3%) was significantly more accurate (t(40) ≥ 5.71, p ≤ 8.16×10^−7^), with LFP power, *I* and *H* classifiers performing similarly to each other (|t(40)| ≤ 1.80, p ≥ .227). The Φ* classifier was better at identifying both effective (F-Score_E_ = 0.79) and control (F-Score_C_ = 0.82) stimulations than the LFP power classifier (Fig. 2, F and G; F-Score_E_ = .63, F-Score_C_ = .68).

Next, we determined how decoding accuracy for LFP power and Φ* classifiers depended on their features. For the LFP power classifier, removing P_d_ or C caused greater MDAs (Fig. 2H) than other areas (P_m_, P_s_, F_d_, F_m_, F_s_, t(280) ≥ 18.35, p ≤ 3.40×10^−15^; Table S1). Removing delta and low/high-gamma frequencies caused significantly greater MDAs (Fig. 2I) than other bands (θ, α, β; t(240) ≥ 4.36, p ≤ 1.39×10^−4^; Table S1). For the Φ* classifier, removing subsystems containing P_d_ or C caused significantly greater MDAs (Fig. 2J) than other areas (P_m_, P_s_, F_d_, F_m_, F_s_, t(140) ≥ 5.96, p ≤ 2.59×10^−7^, Table S1). These results suggest parietal and subcortical contributions to stimulation-induced consciousness.

We next tested how well each measure tracked finer-scale changes in consciousness level induced by thalamic stimulation. We computed correlations between changes in consciousness level and normalized changes (during – pre) in either LFP power (42 features: 7 areas x 6 frequencies), Φ*, *I* or *H* (for full 7-part system). For LFP power (Fig. 3, A-H), *I* and *H* (Fig. S6), correlations were not significant. In contrast, normalized changes in Φ* (Fig. 3I) correlated with consciousness level (r = 0.62, t(160) = 10.33, p < .0001). Linear regression analysis confirmed these results (Fig. 3, B-I; Fig S6; Table S2).

**Fig. 3.**
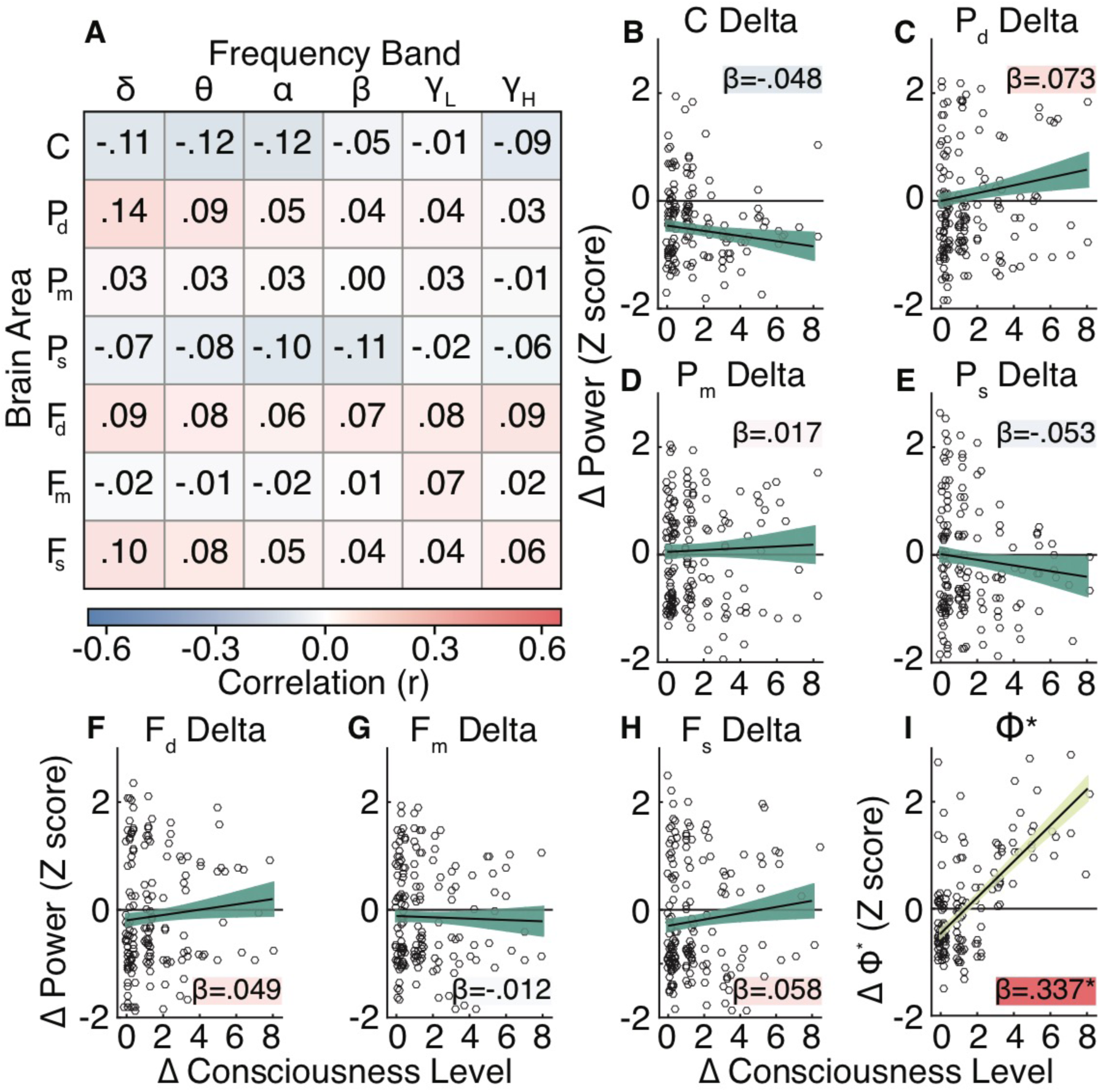
Only Φ* correlates with fine, stimulation-induced changes in consciousness level. (**A**) Correlations (r) of normalized changes in power (Z score, during stim – pre) and changes in consciousness level (during stim – pre) for each brain area at each frequency band. (**B-H**) Regression estimates of delta power correlations in (A) for (**B**) caudate nucleus, (**C**) deep, (**D**) middle and (**E**) superficial parietal layers, as well as (**F**) deep, (**G**) middle and (**H**) superficial frontal layers. Circles show values for each stimulation event. Line shows regression fit (± SE) with reported slope (β) and correlation (shaded background, r, on same scale as (A)). (**I**) Regression estimate for stimulation-induced consciousness level changes (during stim – pre) on Φ*. * on β slope indicates p = 9.0×10^−15^.

Φ* consistently outperformed other measures and had a reliable relationship to even small changes in consciousness, supporting IIT. Within the IIT framework, system parts contributing most to integrated information contribute most to consciousness (Fig. S1D). We compared average Φ* across all subsystems during wakefulness, sleep and anesthesia, as well as prior to, during and after effective/control stimulations, as a function of the composition of each subsystem and consciousness (Fig. 4, A-D). For the state data (Fig. 4A), subsystems containing P_d_, T and C had higher Φ* than systems with other areas (t(378) ≥ 5.90, p ≤ 9.82×10^−8^; Table S3). Effect size estimates showed P_d_, T and C combined (ΔR^2^ = .49), and consciousness level (ΔR^2^ = .39), explained most of the variance in Φ*, whereas frontal cortex only accounted for 4%. Similarly for the stimulation data (Fig. 4C), subsystems with P_d_ and C had higher Φ* (t(372) ≥ 5.55, p ≤ 5.39×10^−7^; Table S3). P_d_ and C (ΔR^2^ = .43), and consciousness level (ΔR^2^ = .29), explained a large portion of variance in Φ*; much more than frontal cortex (ΔR^2^ = .10; Fig. 4D).

**Fig. 4.**
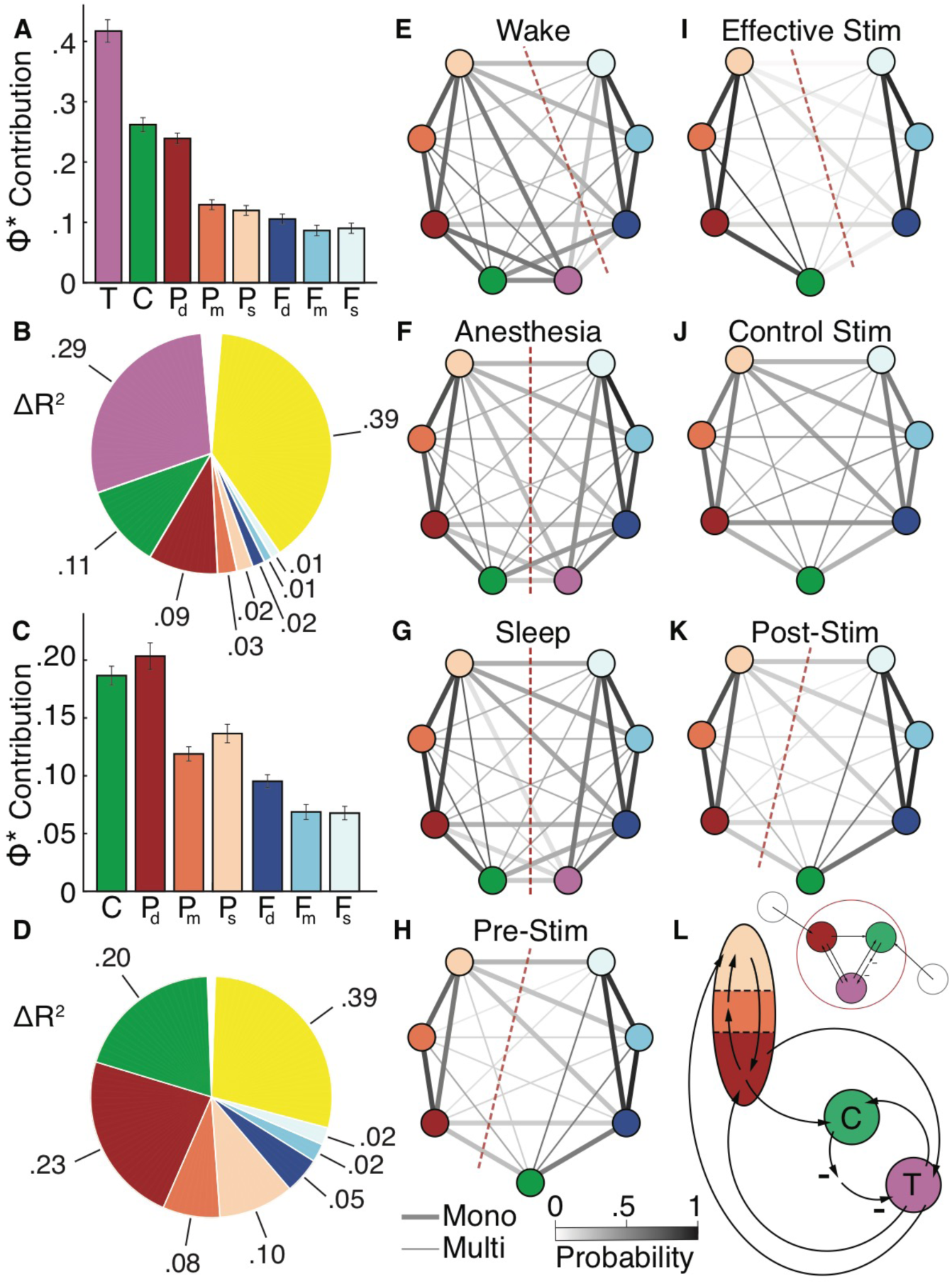
Integration between parietal deep layers and subcortical areas contribute most to increases in Φ* and changes in conscious state. (**A**) Average contribution of each brain area to Φ* (± SE) controlling for changes in conscious state. (**B**) Estimated effect size (ΔR^2^) of each brain area in (A) and consciousness (yellow, wake vs sleep/anesthesia). (**C**) Average contribution of each brain area to Φ* (± SE) controlling for stimulation-induced changes in consciousness. (**D**) Estimated effect size (ΔR^2^) of each brain area in (C) and consciousness (yellow, effective vs pre/post/control stimulations). (**E-K**) Pairwise probability (gray-scale lines) of brain areas associating on the same side of the minimum information partition (MIP) for (**E**) wakefulness, (**F**) anesthesia, (**G**) sleep, (**H**) pre-stimulation, (**I**) effective stimulation, (**J**) control stimulation and (**K**) post-stimulation. Monosynaptic (Mono, thick lines) anatomical paths (in at least one direction) and multisynaptic (Multi, thin lines) shown. Dashed red lines shows predominant MIP when applicable. (**L**) Schematic showing pathways and brain areas contributing most to integration and classification of consciousness.

Calculating Φ* requires determination of the minimum information partition (MIP) for a system, the “cut” that leads to the least amount of information loss (Fig. S2C). Analyzing MIPs thus provides insights about integrated structures across conscious states. We compared kernel density-estimated distributions of the MIP probability mass functions for each state. Sleep (Kolmogorov-Smirnov, KS(42) = .36, p = .0067) and anesthesia (KS(42) = .38, p = .0031) were different from wakefulness (Fig. S7A), but similar to each other (KS(42) = .17, p = .56). The same approach for thalamic stimulations (Fig. S7B) showed effective stimulations (KS(30) = .63, p = 4.64×10^−6^), but not controls (KS(30) = .23, p = .34), had a different MIP distribution from pre-stimulation (similar distributions pre- and post-stimulation, KS(30) = .10, p = .99).

Because MIP distributions differed according to consciousness, we tested which brain areas constituted integrated structures across state and stimulation conditions. We computed the pairwise probability of system parts being on the same side of the MIP (Fig. 4, E-K; Fig S7 F-K). Wakefulness (Fig. 4E) showed increased association of T and P (excluding F), whereas anesthesia (Fig. 4F) and sleep (Fig. 4G) showed increased association of T and F (excluding P; Fig. S7C). Effective thalamic stimulations (Fig. 4I) showed increased association of C and P (excluding F), unlike pre-stimulation and controls (Fig. S7, D and E). Pre- (Fig. 4H) and post- (Fig. 4K) stimulation both showed increased association of C and F (excluding P); while control stimulations (Fig. 4J) had random associations between all areas. Overall, subcortical areas shifted their affiliation from frontal to parietal cortex with increased consciousness. Together, all analyses suggest parietal-striatal-thalamic systems (T, C, P_d_, P_m_, P_s_) are the strongest NCC (Fig. 4L). Further “cuts” of this particular subsystem confirmed MIPs in conscious states favor association of T, C and P_d_ (Fig. S7, F-N). This suggests integration of subcortical areas and parietal deep layers is a hallmark of consciousness (Fig. 4L).

We have shown that parietal deep layers and subcortical areas contributed most to decoding conscious states; whereas frontal cortex, and superficial cortical layers broadly, contributed little, irrespective of metric. These results do not support GNW and HOT, which emphasize frontal cortex (and/or superficial layers), nor RPT, which emphasizes localized cortical processing. Rather, our results are most consistent with IIT, which features parietal contributions and integration. Parietal cortex contains a number of topographic spatial maps in eye, body and world-centered reference frames (*23*). Such spatial representations are vital for perception and awareness, as parietal lesions give rise to spatial neglect (*24*). Based on IIT, grid-like structures, as in parietal cortex, may be especially suitable for supporting our environmental experiences (*25*).

While *H, I* and Φ* all measure complexity, only Φ*, through the partitioning step, measures integration across system parts. As Φ* outperformed other metrics in predicting changes in consciousness, integration – between parietal deep layers and subcortical areas – seems to be vital. However, IIT, with its cortical focus (*10, 16, 25*), has not incorporated subcortical areas like the striatum. Deep layers of parietal cortex project to the striatum and reciprocally connect with thalamus. Further, the striatum and thalamus are famously interconnected by multi-synaptic striato-thalamic pathways, and the often overlooked direct excitatory thalamo-striatal pathway (*17, 18*). These cortico-striatal-thalamic architectures (with multiple feedback loops between C and T, and between T and P) could serve as cause-effect structures proposed by IIT and thus key NCC.

## Acknowledgments

We thank the Tononi and Cirelli Labs, and D. Cleveland, for useful discussions.

## Author contributions

M.R., J.P., N.K., S.M., A.R. and Y.S. performed research; M.A., M.R., A.H. and Y.S. analyzed data; M.A., M.R. and Y.S. wrote paper; M.A., M.R., J.P., N.K., S.M., A.H., A.R. and Y.S. edited paper.

## Data and materials availability

All data and code available upon reasonable request.

## Supplementary Materials

### Materials and Methods

The University of Wisconsin-Madison Institutional Animal Care and Use Committee approved all procedures, which conformed to the National Institutes of Health Guide for the Care and Use of Laboratory Animals. Data relating to the caudate nucleus (C) have not previously been published, but an analysis of much of the frontal eye field (F), lateral intraparietal area (P) and central lateral thalamus (T) can be found in (*22*). Here, however, we focus on new analyses and previously unpublished findings. We acquired data from two male monkeys (*Macaca mulatta*, 4.3-5.5 years old, 7.63-10.30 kg body weight) housed at the Wisconsin National Primate Research Center (WNPRC). Experimenters and WNPRC husbandry staff provided daily animal care, and WNPRC veterinarians monitored animal health.

#### Surgery

We performed a head implant and craniotomy surgery and using aseptic techniques on anesthetized monkeys. We induced anesthesia with ketamine (up to 20 mg/kg body weight, i.m.) and maintained general anesthesia with isoflurane (1-2%). We used 12 ceramic skull screws and dental acrylic to affix a customized plastic recording chamber and head post to the skull. This implant included four hollow slots, two on the left and two on the right sides of the acrylic implant, into which custom designed rods fitted, allowing head immobilization during electrophysiological recordings. We drilled 2.5 mm craniotomies in the frontal and parietal bones within the recording chamber, providing access to F, P, T and C in the right hemisphere. We derived craniotomy coordinates from the high-resolution T1-weighted structural images acquired prior to the surgery, in consultation with a brain atlas (*26*). We fitted each craniotomy with a conical plastic guide tube filled with bone wax, through which linear micro-electrode arrays traversed. We prefabricated these guide tubes using a model of the skull based on the T1-weighted structural image (*13, 22, 27*). We also inserted two titanium skull screws within the recording chamber, one from which to record the EEG and one to serve as a reference.

#### Neuroimaging

We used the GE MR750 3T scanner to perform structural imaging on anesthetized monkeys. At the start of each imaging session, we pre-medicated the monkey with ketamine (up to 20 mg/kg body weight) and atropine sulfate (0.03-0.06 mg/kg). We then intubated the monkey and administered isoflurane (1-2% on ∼1 L/min O_2_ flow), with a semi-open breathing circuit and spontaneous respiration, to maintain general anesthesia for the session duration. We monitored the monkey’s expired carbon dioxide, respiration rate, oxygen saturation, pulse rate and temperature, using an MR-compatible pulse oximeter and rectal thermometer. Prior to the implant and craniotomy surgery, we acquired a high-resolution structural brain image, to delineate the regions of interest (ROIs), F, P, T and C. For these three dimensional T1-weighted structural images, we used an inversion-recovery prepared gradient echo sequence with the following parameters: FOV=128 mm^2^; matrix=256 x 256; no. of slices=166; 0.5 mm isotropic; TR=9.68 ms; TE=4.192 ms; flip angle=12°; inversion time (TI)=450 ms. To generate the high-quality structural image for each monkey, we used NEX = 6-10 T1-weighted structural images, and performed the averaging using the FMRIB Software Library (FSL) (*28*). After surgery, we acquired additional scans of electrodes *in situ* to confirm electrode positioning, using the same set of parameters. To localize implanted electrodes, we used NEX = 2.

#### Electrophysiological recordings

We simultaneously recorded local field potentials (LFPs) from F, P, T and C, using linear micro-electrode arrays (in early C recordings, we used a sharp tip platinum-iridium electrode). There were 16 or 24 electrode contacts positioned in F and P, 24 contacts positioned in T, and 16 (or 1) contacts positioned in C. These platinum/iridium electrode contacts had a diameter of 12.5 μm, 200 μm spacing between contacts, and typical impedance of 0.8-1 MΩ. We measured EEG using titanium skull screws located above dorsal fronto-parietal cortex and, in anesthetized experiments, the EMG using a hypodermic needle (30G) in the forearm. We recorded electrode signals (filtered 0.1-7,500 Hz, amplified and sampled at 40 kHz) using a preamplifier with a high input impedance headstage and OmniPlex data acquisition system controlled by PlexControl software. We recorded wake and sleep data in the same recording sessions, and anesthesia data (with and without thalamic stimulation) in different sessions.

#### Electrode array localization

We targeted electrodes to ROIs based on the T1-weighted structural images of electrodes *in situ* held by the customized guide tubes. Although the electrode itself is not visible in the images, there is a susceptibility “shadow” artifact of approximately one voxel (0.5 mm^3^) width on each side of the electrode shaft. We re-positioned electrodes as necessary, based on volumes acquired online and in reference to a stereotaxic brain atlas (*26*), and re-acquired T1-weighted scans until electrodes were in their desired locations in F, P, T and C. Offline, we used FSL (FLIRT) to register (6 degrees of freedom) the images with electrodes *in situ* to the high-resolution structural image acquired prior to surgery (*29*). We reconstructed recording and stimulation sites along electrode tracks, using measurements of electrode depth during recording sessions calibrated to the electrode depth measurements during imaging sessions and the image of electrodes *in situ*. In one monkey, we also performed post-mortem histology to further verify electrode track reconstructions. We fixed the brain in 10% neutral buffered formalin, and afterwards cut the right hemisphere into approximately 5 mm thick coronal sections. After embedding these sections in paraffin, we thinly sectioned tissue into 8 μm slices. Around ROIs, we stained slices with Hematoxylin and Eosin, and visualized slices under a microscope to confirm electrode tracks through our ROIs.

We further verified recording sites in our ROIs using functional criteria. In the F ROI, we elicited eye movements using local electrical stimulation with low currents, i.e., <100 μA (*30*). In the P ROI during awake experiments, many neurons showed the classical response characteristic of peri-saccadic activity. In the T ROI, we found a subset of neurons with high firing rates (around 40-50 Hz) in the awake state, consistent with a T locus (*31*). To position electrode arrays across cortical layers in F and P, we used depth measurements derived from structural images to initially position arrays in F and P. The 16 and 24 contact arrays respectively covered 3 mm and 4.6 mm (200 μm spacing between electrode contacts), which generally allows for positioning of contacts spanning all cortical layers, for tracks near perpendicular to the cortical surface or with moderate angles from perpendicular. We further adjusted electrode position to maximize the number of contacts showing single-unit or multi-unit spiking activity, and we visualized evoked LFP responses to auditory tones, with middle layers showing earliest response. Offline, we used current source density (CSD) analysis to designate contact positions to superficial, middle and deep cortical layers.

#### Passive auditory oddball

We used a passive auditory oddball paradigm for stimulus-aligned CSD analyses. This allowed identification of superficial (s), middle (m) and deep (d) layers of F and P. The passive auditory oddball paradigm does not require a behavioral response, nor does it require open eyes. Moreover, auditory stimuli have been shown to activate neurons in F (*32-34*) and P (*35-38*). In the oddball paradigm (written in Neurobehavioral Systems Presentation software), we presented a sequence of auditory tones (200 ms duration; 800 ± 100 ms jitter between tones) comprised of 80% standard tones (0.9 kHz) and 20% deviant/oddball tones (1 kHz). At least the first four tones of a sequence were standard tones, and two sequential tones could not be deviants. Otherwise, there was a pseudorandom tone order within the constraint of the overall 80/20 standard-to-deviant ratio. The sequence duration was 3 min for anesthesia experiments, and 6 min for awake experiments. We presented tones through two transducers placed 35 cm from each ear in anesthesia experiments, and 80 cm from each ear during awake experiments. The sound level at each ear was about 75 dB SPL for both anesthesia and awake experiments.

#### CSD

We applied inverse CSD analyses to localize electrode contacts to superficial, middle or deep cortical layers (*39*). We used the CSDplotter toolbox for MATLAB (https://github.com/espenhgn/CSDplotter; dt = 1 ms, cortical conductivity value = 0.4 S/m, diameter = 0.5 mm) to calculate the inverse CSD in response to auditory tones in the passive oddball paradigm. Linear micro-electrode arrays measure the LFP, *v*, at N different cortical depths/electrode contacts along the *z*-axis with spacing *h*. The standard CSD, *c*_*st*_, is estimated from the LFPs using the second spatial derivative, i.e.,

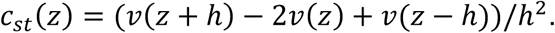

LFPs can also be estimated from given CSDs, represented in matrix form as *V* = ***F****Ĉ*, where *V* is the vector containing the *N* measurements of *v, Ĉ* is the vector containing the estimated CSDs, and ***F*** is an *N* × *N* matrix derived from the electrostatic forward calculation of LFPs from known current sources. The inverse CSD method uses the inverse of ***F*** to estimate the CSD, i.e., *Ĉ* = ***F***^**-1**^***V***. For the step inverse CSD method used here (*39*), it is assumed that the CSD is step-wise constant between electrode contacts, so the sources are extended cylindrical boxes with radius *R* and height *h*. In this case, ***F*** is given by:

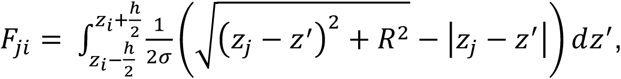

where *σ* is the electrical conductivity tensor, and 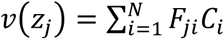 is the potential measured at position *z*_*j*_ at the cylinder center axis due to a cylindrical current box with CSD, *C*_*i*_, around the electrode position *z*_*i*_. The inverse CSD method offers advantages over the standard CSD. The inverse CSD method estimates the CSD around all *N* electrode contacts, whereas the standard CSD method yields estimates around *N* − 2 contacts. Further, the standard CSD requires equidistant contacts, whereas the inverse CSD method does not, which is advantageous when data from a noisy contact may need to be excluded.

We designated the electrode contact at the bottom of the early current sink in response to auditory stimulation (around the boundary between layers 4 and 5) and the two more superficial contacts as the middle cortical layers. We then designated electrode contacts superficial to the middle layers as the superficial layers, and contacts deeper than the middle layers as the deep layers. We cross-referenced layer assignments to the reconstructions of the recording sites along the electrode track, as well as to single-unit and multi-unit spiking activity, which helped delineate the border between gray and white matter. We excluded contacts located outside the ROI from data analyses. Previous studies generated CSD data in F (*40*) and P (*41*) using visual stimulation. Consistent with these studies, our auditory-aligned CSD data also showed sensory stimulation eliciting early sinks in middle layers. In addition, we performed CSD analyses using LFP signals aligned to the trough of delta-band oscillations recorded from the electrode contact with the highest delta power, i.e., this contact served as the phase index (*42-44*). These delta phase-realigned CSDs showed differences across cortical layers, which helped verify that electrode positions remained stable across resting state recording blocks (in which there is no auditory stimulation).

#### Behavior

To match behavioral and sensory conditions across different levels of consciousness, we acquired electrophysiological data from monkeys in a dark, quiet room during wakeful resting state (no task requirements/no report condition), light non-REM sleep, general anesthesia, and stimulation-induced arousal from general anesthesia. To additionally control for eye movements in awake monkeys, we also acquired electrophysiology data during a fixation task (written in Neurobehavioral systems Presentation software), in which the monkey needed to fixate a central dim gray circle of diameter 0.42° visual angle on a black monitor screen located 57 cm away. The monkey received 0.18-0.22 mL of juice every 2.2-3.5 s while maintaining fixation within a window of 3°x3° visual angle, centered on the dim gray circle. When the monkey’s gaze left the fixation window, he typically re-established central fixation quickly. We doubled the juice volume when fixation persisted beyond 10 s, which encouraged long fixations. For awake experiments, we monitored the monkey’s eye position using a video-based eye tracker with 500 Hz sampling rate. For anesthesia experiments, we monitored eyes using a digital video camera capturing 30 frames per second. We measured luminance contrast in a window tightly bounding the eye image using custom-written software in Matlab. This contrast measure differentiated closed eyes, i.e., relatively homogenous high luminance eyelid shade, from stimulation-induced eye openings, i.e., dark pupil and iris contrasting against sclera. Visual inspection of the eye video verified the timing of eye openings/closings derived from the contrast measure.

#### Awake experiments

There were 40 awake experimental sessions (18 for monkey R; 22 for monkey W), each usually 2-4 hours in duration. Monkeys sat upright in a primate chair with their head immobilized using the head post and/or four rods that slid into the hollow slots of the acrylic implant. Awake experiments were split into two types, similar to the two phases of anesthesia experiments. The first type involved simultaneous recordings from F, P, T and C; and the second type involved T stimulation while recording from F, P and C. For both types of experiments, we performed recordings across multiple blocks of resting state, the fixation task and passive auditory oddball paradigm, interleaving blocks involving reward (fixation task) with those not involving rewards (resting state and passive oddball). We randomly varied the specific order across different experimental sessions. Our ROIs have been implicated in awareness, but they also contribute to selective attention and oculomotor function. Thus, we controlled for effects related to attentional or saccadic processes. For awake experiments, we compared recordings during resting state to the fixation task. We only analyzed electrophysiological data during stable eye epochs, i.e., when eye position remained fixed for at least 1 s. This applied to all wake-state data in the resting state and fixation task. Because the wakeful resting state and fixation task data were similar, they were combined for the wake dataset.

#### Sleep

In awake experiments, monkeys at times would fall asleep, particularly during resting state. Online, we identified non-rapid eye movement (NREM) sleep using the following criteria: (i) high EEG delta (1-4 Hz) activity; (ii) extended eye closure (we recorded times when eyes closed and re-opened); (iii) preceding period of drowsiness indicated by drooping/closing of eyelids; (iv) disengagement from fixation task (if applicable); and (v) musculoskeletal stillness. Offline, we identified NREM sleep periods using EEG and eye tracker data. We bandpass filtered EEG data (1-4 Hz; Butterworth, order 6) and applied the Hilbert transform, to calculate the instantaneous delta-band amplitude. From the resulting time series, we detected times of relatively high delta amplitude using thresholds titrated for each experimental session, because the mean delta amplitude and standard deviation could vary depending on the session and total sleep time. For each session, we selected the threshold as the number of standard deviations from the mean delta amplitude that produced a total sleep time estimate that closely resembled the expected sleep time based on online NREM sleep identification, as well as the offline calculation of the total time when the monkey’s eyes were closed, using the recorded eye tracker time series data. Offline NREM sleep identification and time stamping then involved automated detection of extended epochs across the session when both the monkey’s eyes were closed, and delta amplitude was above threshold. These offline NREM sleep detections were consistent with online detections and proved reliable across sessions and monkeys. The identified NREM sleep epochs corresponded to N1 or N2 phases, i.e., light sleep. Thus, monkeys were at a different depth of unconsciousness during sleep compared to general anesthesia in our study. The LFP data during sleep allowed us to compare the influence of conscious and less-conscious states on the same subset of LFPs recorded in both wakefulness and sleep. This further substantiated our comparison between the awake and anesthetized states, in which we used two different subsets of LFPs from the same ROIs, because maintenance of stable anesthesia up to 12 hours required recordings to occur in a surgical suite, whereas awake experiments took place in the behavioral lab.

#### Anesthesia experiments

We used two different anesthetic agents, either propofol (9 sessions: 4 for Monkey R, 5 for Monkey W) or isoflurane (9 sessions: 5 for Monkey R, 4 for Monkey W), to ensure that results reflected general mechanisms of anesthesia/consciousness, and not drug-specific effects. Each anesthesia experimental session was 10-12 hours in duration. After inducing anesthesia with ketamine (up to 20 mg/kg body weight, i.m.), we intubated the monkey and inserted an intravenous catheter(s) for fluid and drug administration. We maintained general anesthesia in spontaneously respiring monkeys with propofol (0.17-0.33 mg/kg/min i.v.) or isoflurane (0.8-1.5% on 1 L/min O_2_ flow), and a clinical anesthesiologist (A.R.) oversaw stable conditions throughout. We stabilized the head of the monkey (in prone position) using four rods that fit into the hollow slots of the acrylic implant. The rods were attached to a modified stereotaxic apparatus atop a surgical table. We maintained the monkey’s temperature using a forced-air warming system and monitored end tidal carbon dioxide, respiration rate, oxygen saturation, heart rate, blood pressure and rectal temperature.

In the first phase of each experimental session, we simultaneously recorded from F, P, T and C. We independently positioned the linear micro-electrode arrays in each ROI, using microdrives coupled to an adapter system allowing flexibility in approach angles across arrays. These recordings started at least two hours after anesthetic induction, and 30 minutes after positioning arrays to allow for settling of tissue. We used a number of different anesthetic levels, adapting the dose to reflect a range of clinically relevant anesthetic depths, e.g., 0.2, 0.225, 0.25 and/or 0.3 mg/kg/min propofol, or 1%, 1.1%, 1.25% and/or 1.5% isoflurane, allowing dosing changes to stabilize before starting the next block of recordings (typically at least 30 minutes). In the second phase of each session, we electrically stimulated T while recording from F, P and C, without changing the anesthetic regimen. For stimulations, we used the linear electrode array existing in T or replaced it with another array inserted along the same trajectory to the same depth. If the first stimulation site did not induce a change in the level of consciousness, then we moved the stimulating electrode to a new depth in the thalamus, in steps of 0.5-1 mm dorsal or ventral along the electrode track, until stimulation induced a change in consciousness level. In early experiments, we tested T stimulations at different anesthetic doses between 0.17-0.3 mg/kg/min for propofol and between 0.8-1.3% for isoflurane. We observed stimulation-induced changes in the level of consciousness for all but the highest doses, i.e., 0.3 mg/kg/min propofol and 1.3% isoflurane. In subsequent experiments, we used propofol doses between 0.17-0.28 mg/kg/min (M = 0.23, SD = 0.03), and isoflurane doses between 0.8-1.25% (M = 1.04, SD = 0.11) during T stimulation. As previously reported (*22*), consciousness level changes did not depend on the anesthetic used, nor the dose. For both experimental phases (recordings with or without stimulation), we interleaved resting state epochs and the passive auditory oddball paradigm.

#### Electrical stimulations

We electrically stimulated T using 24-contact electrode arrays, which had previously been used several times as recording electrodes and thus had lower impedance. We simultaneously stimulated via 16 electrode contacts, with 400 μs bi-phasic pulses of 200 μA, at 50Hz frequency, for a total of 60 s stimulation duration for each stimulation event (experiments included multiple stimulation events). This stimulation protocol has been shown to reliably increase the consciousness level in (the same) anesthetized monkeys; see (*22*) for further validation of electrical stimulation methods and behavioral effects. We typically performed three stimulation events within a stimulation block for reproducibility, with a recovery time of at least the stimulation event duration between repetitions, i.e., stimulations from 1-2 minutes, 3-4 minutes, and 5-6 minutes of a seven minute block.

#### Scoring the level of consciousness

We quantified the level of consciousness induced by thalamic stimulation, using a customized index incorporating five indicators, each scored 0, 1 or 2. The sum of the scores of the five indicators yielded a consciousness level index, ranging from 0-10. The five indicators are:

1. limb/face movements (0 = nothing; 1 = small movement or increased EMG with no clear movement; 2 = full reach or withdrawal)
2. oral signs (0 = nothing; 1 = small mouth/jaw/tongue movements; 2 = full jaw openings/closures, with multiple repetitions)
3. body movements (0 = nothing; 1 = small torso movement or swallowing; 2 = large full torso movement)
4. eye movements/openings (0 = nothing; 1 = eyelid flutters/small blinks or increased eye movements; 2 = full eye opening with occasional blinks)
5. vital signs (0 = no change, i.e., difference of <10% respiration rate (RR), <5% heart rate (HR); 1 = difference of >10% RR, >5% HR; 2 = at least 20% change in either RR or HR, or at least 10% change in both RR and HR; compared to baseline 30 s prior to stimulation).

A WNPRC veterinarian, a clinical anesthesiologist, and five other primate electrophysiologists observed the electrical stimulation effects during anesthesia experiments. We scored the consciousness level before, during and after all stimulation events, using observations recorded during experiments, as well as offline review of videos and EMG data (filtered 30-450 Hz, full-wave rectified, then filtered 5-100 Hz to extract the envelope). A typical stimulation block consisted of three stimulation event repetitions (one minute each) within a seven minute period at a given site, using the same stimulation frequency, current, polarity, duration, anesthetic and dose. We defined stimulation event epochs from the onset to offset of stimulation pulses, i.e., from 1-2 minutes, 3-4 minutes, and 5-6 minutes of a seven minute block. The time between two stimulation epochs was split equally into post- and pre-stimulation epochs. The pre-, during and post-stimulation index of consciousness level for a block reflected the maximum possible score across the repetitions (repetitions largely produced the same score within each epoch type). Prior to electrical stimulations, the consciousness level index was 0 or 1. This could be differentiated from stimulation events inducing an index of 3 or more by all observers. Thus, we defined effective stimulation events as those inducing an index ≥3 (typically stimulation centered on T), whereas control (ineffective) stimulation events had an index of 0-2 (typically stimulation not centered on T).

#### Preprocessing

We divided electrophysiological data time series into epochs of 1 s duration for analysis. In the anesthetized and non-REM sleep states, we divided LFPs into non-overlapping 1 s epochs. In the awake state, we first identified time periods when the eyes were stable, i.e., periods starting 200 ms after a saccade and ending 200 ms before the next saccade. Next, we divided stable eye periods into non-overlapping 1 s epochs. This matched oculomotor state (stable eye position) in data epochs from conscious and unconscious states.

We lowpass filtered data below 250 Hz for LFPs (Butterworth, order 6, zero-phase filter) and down-sampled to 1 kHz. Next, we linearly detrended LFPs, and extracted artifacts (power line noise) by removing significant sine waves using the Chronux (http://chronux.org/) function rmlinesc. We then calculated bipolar derivations of LFPs, i.e., the difference between two adjacent electrode contacts (excluding contacts that had been removed due to noise), to minimize any possible effects of a common reference and volume conduction (*43-45*).

During electrical stimulations, there was a brief artifact caused by the applied current. To remove this artifact, we first excised a 1 ms window around the artifact, then linearly interpolated across this window. Next, we used the Chronux function rmlinesc to remove any significant sine waves at the stimulation frequency (we also performed artifact removal using the SARGE toolbox (*46*), which yielded qualitatively similar results).

We excluded any data epochs in which bipolar-derived LFPs from F, P, T or C had amplitude greater than 3.5 standard deviations from the mean, for both information theoretic and LFP power calculations. Prior to entropy *(H)*, mutual information *(I)* and Φ* calculations, we normalized each LFP based on its mean across all epochs for that recording session. Finally, each LFP was binarized with respect to its median amplitude over the 1 s epoch, removing potential biases related to amplitude differences across channels or conditions.

#### Calculating Φ*

Due to the complexity of the system from which we recorded and its states at each time point, calculating integrated information (Φ) directly is intractable (*47*). However, with certain assumptions about the organization of included brain areas, integrated information can be estimated from the measure Φ* (*20, 47, 48*) (Fig. S2, A-C). To compute Φ*, we construct the state of a subsystem *X*(*t*) at time *t* (1 ms time bins considering 1 kHz sampling frequency) as:

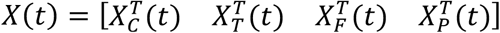

where its elements are the bipolar-derived LFP signals, ranging from 1 to 23 signals (derived from 24 electrode contacts) for each area, C, T, F and P. That is, the component of *X*(*t*) for each brain area is *N*_ch_ × *T*-sized, where *N*_ch_ specifies the number of bipolar-derived channels for each one of four areas and *T* = 1000 (1 s epochs, sampled at 1 kHz). Next, we calculate the uncertainty about the states assuming a multivariate Gaussian distribution of the states across time as:

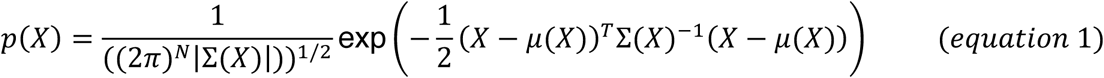

where Σ(*X*) (*t* is removed for brevity) is the covariance matrix of *X* estimated over a 1 s data epoch, *σ*_*ij*_ is the covariance between channels *i* and *j* of the state vector *X* over the epoch, |Σ(*X*)| is the determinant of the covariance matrix that can be considered as a measure of uncertainty about the state *X* at any time point within the epoch, and *N* is the total number of channels across areas. We computed the relationship between the ordered Mahalanobis distances of the data from the mean matrix and their corresponding χ^2^ quantiles, to ensure the states have a Gaussian distribution. The entropy for the states *X*(*t*), given its probability density function (pdf) as *p*(*X*), is defined as:

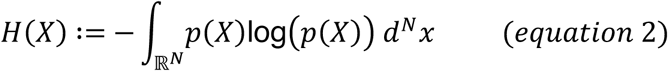

The entropy will be maximized if *p*(*X*) is a multivariate Gaussian, and can be calculated in closed form as:

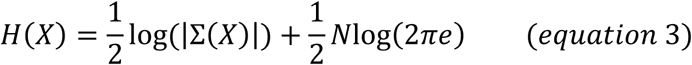

which can be described as the uncertainty about the state *X*(*t*) at time *t* (Fig. S2C). The reduction of uncertainty about the state *X*(*t*) at time *t*, given its past at time *t* − *τ*, also known as the mutual information, can be calculated as:

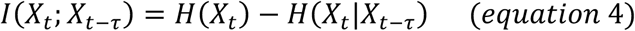

where *H*(*X*_t_|*X*_*t*−*τ*_) is the conditional entropy of state *X*(*t*) given its past state *X*(*t* − *τ*), and can be derived in canonical form with a Gaussian distribution assumption of states as:

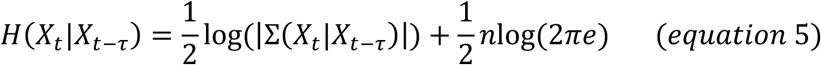

where Σ(*X*_t_|*X*_*t*−*τ*_) is the covariance matrix of the conditional distribution *p*(*X*_t_|*X*_*t*−*τ*_) (conditional covariance) that can be expressed analytically as:

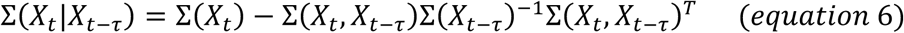

Mutual information, *I*, can be considered a measure of the information the current state has about its past, and it is used to calculate Φ*, a measure of integrated information (Fig. S2D). Φ* of the subsystem *X*(*t*) is information that cannot be partitioned into independent parts of *X* and can be defined as:

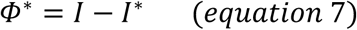

where *I*\* (disconnected *I*) is called mismatched information (Fig. S2C). *I*\* was calculated for every bipartition of system *X*. For most conditions 8 system parts were defined: (C, T, F_s_, F_m_, F_d_, P_s_, P_m_, P_d;_ where s, m and d subscripts correspond to superficial, middle and deep layers respectively). From this, 247 subsystems can be tested (28 two-channel subsystems, 56 three-channel subsystems, etc). Within each cortical layer (F_s_, F_m_, etc) there were multiple bipolar-derived channels; these were never partitioned, and so each layer was effectively a single subsystem part consisting of multiple parallel channels.

The partition *P* that minimizes the normalized Φ* is the *minimum information partition* (*MIP*), as defined in (*49*):

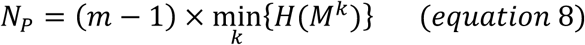

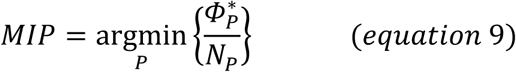

Here *m* is the number of partitions and *M*^*k*^ is the *k*^*th*^ part of subsystem *X*. N_P_ counterbalances inevitable asymmetries introduced by computing *Φ*\* across a variable number of partitions of unequal sizes. The *MIP* reveals the weakest link between the parts of *X*, where the information loss by the partition into subsystems is minimal. We calculated the covariance and cross covariance matrices for each 1 s epoch using the shrinkage method for a more stable result, and averaged them across all epochs for each recording session to calculate *MIP* and its corresponding *Φ*\* 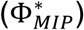, using gradient ascent and L-BFGS optimization method. We incorporated 2 NVIDIA GTX 1080ti GPUs to speed-up the process of searching for MIP to calculate 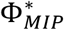

We initially calculated *I* and Φ* for 8 discrete lag times (*τ* ∈ {5, 10, 15, 20, 30, 40, 50, 100 ms}) to find the optimum value for our analyses. We selected *τ* = 15 *ms* for all further analyses, because it yielded the highest Φ* for all conditions. Of note, this *τ* is in the gamma frequency range, which, in parallel analyses, proved consistently important for decoding conscious state based on spectral power. We calculated Φ* for all non-trivial and unique subsystems. For the state data (wake, sleep, anesthetized), there were 8 system parts (C, T, F_s_, F_m_, F_d_, P_s_, P_m_,, P_d_), giving rise to 247 non-trivial subsystems. For the thalamic stimulation data in anesthetized monkeys (pre, during, post-stimulation), thalamic recordings were not possible, thus there were 7 system parts (C, F_s_, F_m_, F_d_, P_s_, P_m_, P_d_), giving rise to 120 subsystems for which we calculated Φ*. For instance, considering the state data without stimulation, there was one system with 8 parts (*s* := {*C*|T|*F*_s_|*F*_m_|*F*_d_|*P*_s_|*P*_m_|*P*_d_}), 8 subsystems with 7 parts, which are derived by removing one of the parts at each permutation (e.g., removing C yields the subsystem *s* := {T|*F*_*s*_|*F*_*m*_|*F*_*d*_|*P*_*s*_|*P*_*m*_|*P*_*d*_}), and so on. We calculated *H* and *I* for the same composition of subsystems (Fig. 1B and Fig. 2,C and D for Φ*; Fig. S3, A-F, for *I* and *H*).

#### Decoding the state of consciousness based on LFP power

To decode the state of consciousness, we calculated the power spectral density (PSD) of bipolar-derived LFPs for each 1 s epoch using the multi-taper method (5 Slepian tapers). Averaging across frequencies of interest and power values within brain areas yielded the mean power for each frequency band (delta 1-4 Hz; theta, 4-8 Hz; alpha, 8-15 Hz; beta, 15-30 Hz; low gamma, 30-60 Hz; high gamma, 60-90 Hz) in each system part (C, T, F_s_, F_m_, F_d_, P_s_, P_m_, P_d_). This generated 48 input features (8 system parts x 6 frequency bands) for the conscious state classification model.

We trained a Support Vector Machine (SVM) model (*50*) to classify the LFP power data into three states: wake (W), sleep (S), and anesthesia (A). We specify *y*_i_(*t*) ∈ {0,1,2} as the identifier of the state of each epoch, where 0, 1, and 2 denote W, S and A respectively. The SVM classifier was implemented by a nonlinear projection of the training data **x** feature space 𝒳 into a high dimensional feature space ℱ using a kernel function *ϕ*. With *ϕ*: 𝒳 → ℱ being the mapping kernel, the weight vector **w** can be expressed as a linear combination of the training inputs (60% of the whole shuffled and stratified input in our model), and the kernel trick used to express the discriminant function as:

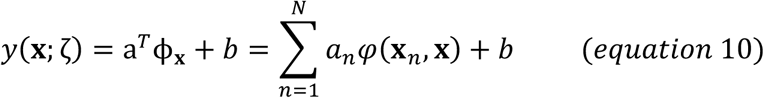

where ζ = {*a, b*} is the new parameter with *a* and *b* as weights and biases of the mapped feature space ℱ respectively. We used the radial basis function (RBF) kernel that allows nonlinear decision boundary implementation in the input space. The RBF kernel holds the elements:

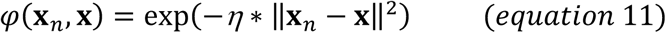

where *η* is a tunable parameter at each cross-validation fold. Model hypermeters consisting of regularization penalty (*C*) and *η* were selected by grid search through 20-fold cross-validation, during which the model accuracy was calculated. To account for the class imbalance in the training and test set, at each cross-validation fold the shuffled stratified dataset was selected as the input to the model.

We used two measures to compare classifiers: accuracy and F-score. Accuracy is the proportion of correctly classified inputs and is defined as:

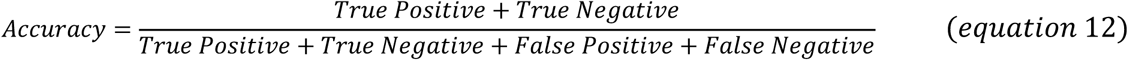

F-score balances classifiers’ precision and recall and is calculated as:

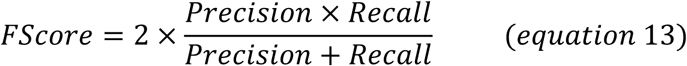

where

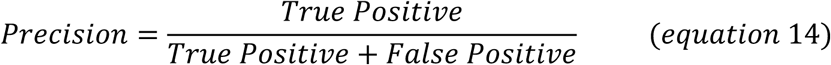

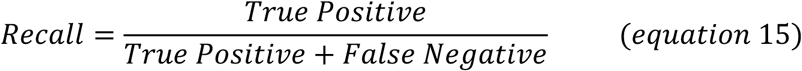

To determine the importance of each input feature, we calculated the mean decrease in accuracy (MDA) after removing the aforementioned input feature from the feature set, then training and performing 20-fold cross-validation of the model again. For instance, to measure the MDA for the T part of the system, the T power for all 6 frequency bands were excluded from the input feature set to the classifier; and to measure the MDA for the delta-band, the delta-band power for all 8 system parts were excluded from the input features.

Before interpreting feature importance, we examined non-independence across features in our classifier by comparing inter-feature correlations (Fig. S5A). Features in the LFP classifier could relate by either brain area or frequency. As expected, correlations were much higher for related features (diagonal) with similar brain areas, than less related features (different areas). To quantify this, we created pools of correlations between features that shared the same brain area or frequency and those that did not, and computed unpaired t-tests. Tests were significant both for brain area (t(924) ≥ 8.20, n_1_ = 21, n_2_ = 903, p ≤ 2.30×10^−4^) and frequency (t(856) ≥ 11.70, n_1_ = 36, n_2_ = 820, p ≤ 8.10×10^−7^). This confirmed that dissimilar features (bound by different areas or frequencies) were not clearly dependent on each other, for each area and frequency band. Because dependence is low and we are cross validating the classifier 20 times to minimize random variance, the MDA approach is a valid measure of feature importance and can be interpreted as the contribution of each system part or frequency band to conscious state decoding.

#### Decoding the effectiveness of thalamic stimulation based on LFP power

We used a similar SVM classifier to decode the effectiveness of thalamic stimulation. LFP power features were extracted for 7 areas and 6 frequency bands, using the same process described in the previous section. We specify *y*_i_(*t*) ∈ {0,1} as the identifier of the effectiveness of thalamic stimulation for that particular epoch, where 0 and 1 denote effective and control stimulations respectively (epochs with stimulation-induced consciousness level index equal to or greater than 3 considered effective; control (ineffective) otherwise). We used the same model and kernel as for the conscious state decoder, except for the hyperparameters which were tuned during model training and 20-fold cross-validation.

#### Decoding the state of consciousness based on Φ*/*I*/*H*

As stated above, we computed our complexity measures from a system of 8 possible brain areas/system parts, giving rise to 247 non-trivial possible subsystems, each of which could serve as a feature for classification. To limit the amount of potentially redundant features, ensure similar conditions (“leave-one-out” for MDA analyses), and better match the number of features for the Φ* and LFP power classifiers, we used <u>Φ*/</u>*<u>I</u>*<u>/</u>*<u>H</u>* for subsystems with 7 or greater number of system parts to classify the state of consciousness. This generated 9 input features for each of Φ*, *I* and *H* – one feature reflects the entire system of 8 parts (C, T, F_s_, F_m_, F_d_, P_s_, P_m_,, P_d_) and the other features reflect all possible combinations of 7 parts. We used the same model, parameters and methods as those used for the LFP power classifier. We trained a model for each feature set separately (Φ*, *I* or *H*) and measured the accuracy for each cross-validation fold.

To further calculate each area’s contribution to classifying the state of consciousness based on Φ*, we trained another model using Φ* values of the subsystems with 6 or greater number of system parts – generating 37 input features – as well as a model using Φ* values of the subsystems without the part of interest – generating 8 features. We then calculated the MDA for the model with the reduced feature set.

Before interpreting feature importance, we examined dependence across features in our classifier by comparing inter-feature correlations (Fig. S5B). Features in the Φ*classifier overlap by subsystem composition. As expected, correlations were much higher for related features (diagonal) all missing the same brain area, than less related features (missing different areas). To quantify this, we used the same method as that used for the LFP power classifier (comparing pools of correlations between features that shared the same brain area and those that did not), and found the correlations to be significantly different (t(471) ≥ 3.45, n_1_ =36, n_2_ = 435, p ≤ .038). Once again, this justified our use of the MDA approach to measure feature importance.

#### Decoding the effectiveness of thalamic stimulation based on Φ*/*I*/*H*

We used Φ*/*I*/*H* for subsystems with 6 or greater system parts to classify the effectiveness of thalamic stimulation. This generated 8 input features for each one of Φ*, *I* and *H* – one feature reflects the entire system of 7 parts (*C, F*_*s*_, *F*_*m*_, *F*_*d*_, *P*_*s*_, *P*_*m*_,, *P*_*d*_) and the other features reflect all possible combinations of 6 parts. We used the same model, parameters and methods as that used for the LFP power classifier for stimulation effectiveness. We trained a model for each feature set separately (Φ*, *I* or *H*) and calculated the accuracy for each one of 20 cross-validation folds. We used the same approach as in the previous section to find each area’s contribution to the decoding of stimulation effectiveness.

#### MIP distribution

The MIP for each recording session can be considered the natural partitioning of a system such that its information is above and beyond its constituent subsystems, as well as the weakest link of the system. Thus, system parts in one partition have a stronger connection among themselves compared to the rest of the system. We used this concept, as well as the frequency of all the bipartition sets for Φ*, to compare how the structure of the system changes across conscious states and thalamic stimulations. We generated histograms of the MIP distribution normalized for each state or stimulation separately, resulting in a probability mass function (PMF) for that condition (Fig. S7, A and B). To estimate distributions of MIPs, we used a gaussian kernel to fit a distribution to each histogram, assuming that the MIP samples are independently and identically distributed as well as drawn from a categorical distribution. We also computed the pairwise probability of each brain area occurring on the same side of the MIP, derived from the frequency of appearance for each MIP type in the PMF. We interpreted these probabilities as connection weights for ease of presentation (Fig. 4, E-K and Fig. 7, F-K).

#### Statistical analyses

Statistical analyses were performed using correlations and linear mixed effect models in R and the Pymc3 (*51*) package in Python. All p-values were controlled using the Holm-Bonferroni method. Where appropriate, models were analyzed using linear models (LM in R), reporting Beta slopes, t statistics, corrected p-values, and ΔR^2^ effect sizes. When necessary to control for other sources of non-independence, we used linear mixed effect models (LMER in R), reporting Beta slopes, F statistics, and corrected p-values. Because it is not a straightforward process to compute ΔR^2^ effect sizes for LMER models, we estimated effect sizes by computing similar LM models without the random effects structure. Given that these models yielded highly similar Beta parameters, their effect sizes serve as reasonable estimates for the more appropriate LMER models.

#### State and Stimulation changes in LFP power, I, H and Φ*

To compare the effects of state on LFP power, we regressed normalized power for each brain area in the delta band on state (wake, sleep, anesthesia) as a dummy coded variable. P-values were corrected for all LFP power tests as a family. So as not to overinflate the sample size, for *I, H and Φ*^*\**^, we averaged values across all possible subsystems for each state sample and then computed the same regression. Tests were corrected for all complexity measures as a family.

To compare the specificity of stimulation effects on LFP power, we regressed normalized delta power for each brain area on stimulation state (during stim = .5, pre/post = −.5), effectiveness (effective = .5, control = −.5), and the interaction. Effects were visually similar at other frequency bands, and so tests were not repeated. Again, to not inflate sample size, for *I, H and Φ*\* we averaged values across all possible subsystems for each stimulation event before computing the interaction.

#### Inter-feature correlation differences

We created pools of similar features (sharing the same brain area or system structure) and different features, then computed unpaired t-tests.

#### Decoding effects

Classifier performances were compared for cross validated repetitions and tested against chance (.33% for state classifiers, .50% for stimulation classifiers). We tested performance between classifiers in a pairwise manner using linear regression and dummy coding with corrected p-values. Mean decreases in accuracy (MDA) were similarly analyzed using dummy coding separately for the LFP power classifier, for brain areas and frequency bands, and for brain areas in the Φ* classifier (see Table S1 for detailed statistics); p-values were corrected within each model feature type.

#### Correlations with stimulation-induced changes in level of consciousness

LFP power for all areas in all frequency bands (δ, θ, α, β, γ_L_, γ_H_), *I, H* and Φ* were normalized within animal and anesthetic. Correlations were computed between changes in each normalized metric (during stim – pre) and the corresponding changes in level of consciousness (during stim – pre). Correlations were computed using the “cor” function yielding r values and T-statistics. Regression estimates were further computed using linear models yielding β slopes and regression lines (Fig. 3, B-I; Fig S6, A and B; see Table S2 for detailed statistics). Covariates for animal and anesthetic were included. No effects were significant for the LFP power data even without controlling for multiple comparisons. Complexity measures (*I, H* and Φ*) were controlled for multiple comparisons as a family.

#### Φ* analyses

To test the contribution of each brain area to Φ*, we regressed average Φ* for each possible system (contained at least 2 brain areas) on consciousness and on variables representing the binary membership (0 = not member, 1 = member) of each brain area in the system. For example, a system containing TP_d_P_m_F_d_F_s_ would be represented as (A_1_:T = 1, A_2_:C = 0, A_3_:P_d_ = 1, A_4_:P_m_ = 1, A_5_:P_s_ = 0, A_6_:F_d_ = 1, A_7_:F_m_ = 0, A_8_:F_s_ = 1). Membership effects were computed within systems of the same size (number of total parts in the system, e.g., the system above has a size of 5). For the state data, consciousness was coded as a state-based contrast (wake = .5, sleep/anesthesia = −.5).

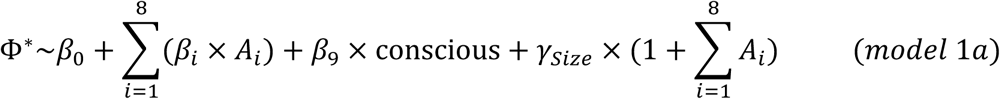

For the stimulation data, we used a similar model that did not contain the thalamus. For this model, consciousness was coded as a contrast of stimulation condition and effectiveness (effective stim = .5, control stim/pre/post = −.5).

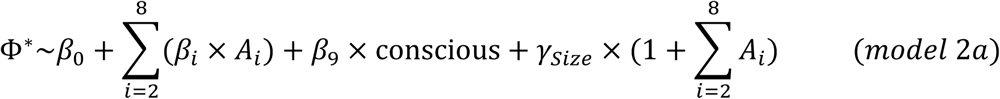

To estimate the effect size of each brain area and consciousness, the same model was computed without the random effect structure. Because the significance and β values were similar, we computed ΔR^2^ for each effect in the model and report this as an estimate of the effect size for each component (Fig. 4, B and D).

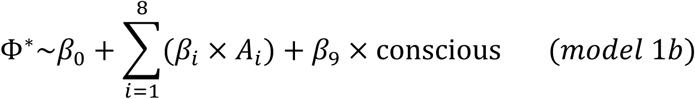

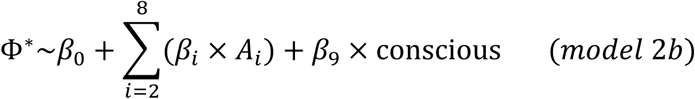

To compare the effects of each area, we computed T-tests between Φ* estimates for systems containing brain area A but not B, and systems that contain area B but not A, for every possible brain area pair and controlled for multiple comparisons (see Table S3).

#### Comparing the probability mass function of MIPs

We applied a two-sample Kolmogorov-Smirnov test using the Pymc3 package in Python, to compare probability mass functions relating to the MIPs across conscious states and stimulations.

---

**Fig. S1.**
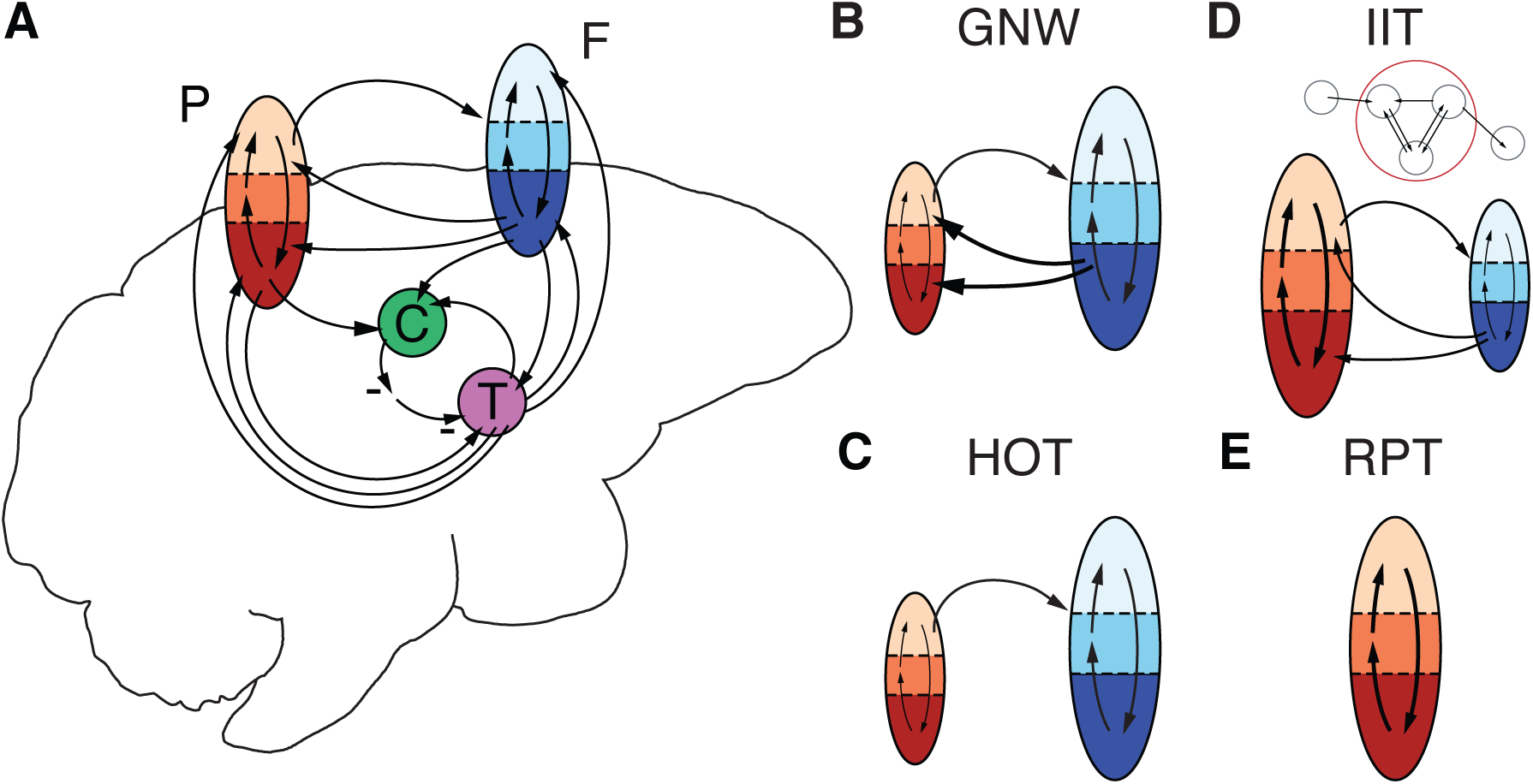
Theories of consciousness are cortico-centric and differentially emphasize contributions of frontal and more posterior brain areas. (**A**) Schematic of anatomical pathways between frontal cortical (F), posterior cortical (parietal, P, or sensory), central lateral thalamic (T) and striatal (caudate nucleus, C) regions of interest in our study. Low, moderate and high color saturation in cortical areas indicates superficial, middle and deep cortical layers, respectively. (**B-E**) Proposed neural correlates of consciousness as postulated by prominent theories, with key brain areas (enlarged) and anatomical pathways (in bold) emphasized: (**B**) Global Neuronal Workspace (GNW); (**C**) Higher-Order Theories (HOT); (**D**) Integrated Information Theory (IIT), with inset showing example mechanism with integrated causes and effects (red circle) generating integrated information; and (**E**) Recurrent Processing Theory (RPT).

**Fig. S2.**
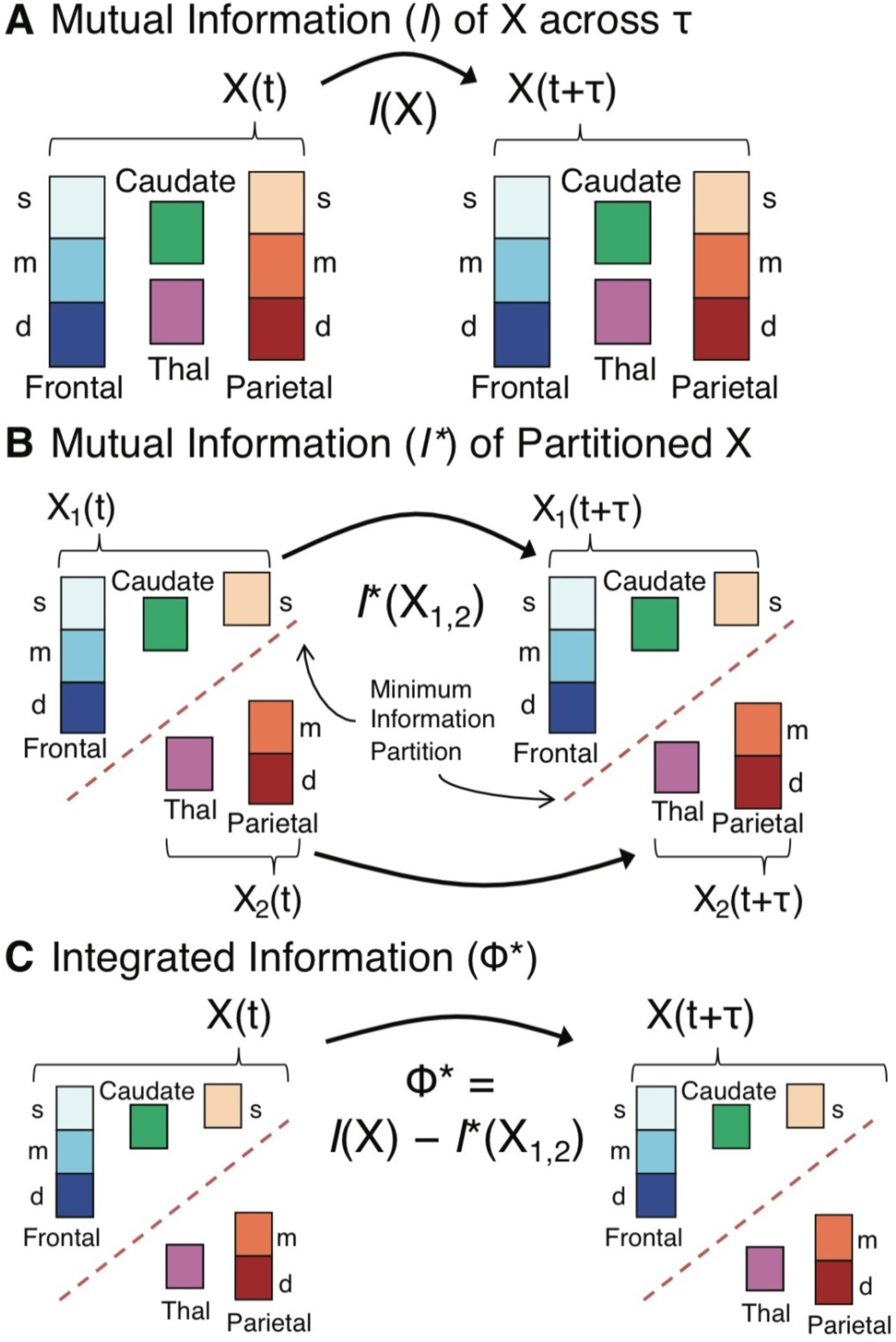
Using mutual information to calculate Φ*. (**A**) Schematic showing calculation of mutual information (*I*) for the full system containing superficial (s), middle (m) and deep (d) layers of frontal and parietal cortex, thalamus (Thal) and caudate nucleus. (**B**) Schematic showing calculation of *I* for a theoretical minimum information partition (*I*^*\**^) represented as a “cut” in the full system (dashed black line). (**C**) Integrated information (Φ*) calculated from *I* and *I*\* for the partitioned system in B. Note that integrated information (Φ) is the metric proposed by Integrated Information Theory to measure consciousness, and Φ* is a computable estimate of that value.

**Fig. S3.**
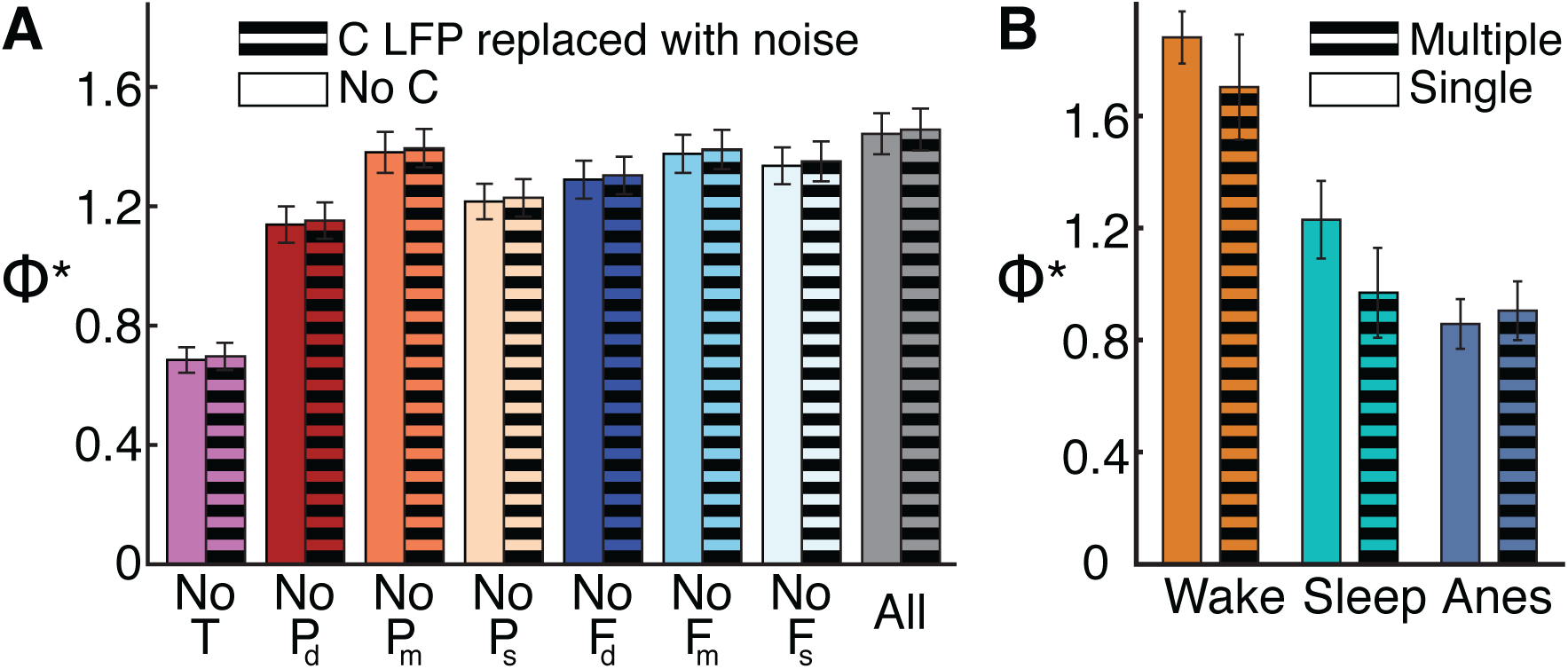
Φ* calculations are neither influenced by noise, nor differences in the number of electrode contacts in each brain area. We tested the (**A**) impact of noise on Φ* by removing the caudate nucleus (C) from the system and adding noise, and (**B**) the impact of the number of electrode contacts on Φ* by comparing single versus multiple contacts in C. (**A**) Φ* (± SE) calculated for different subsystems absent thalamus (T), superficial (s), middle (m) or deep (d) layers of parietal (P) or frontal (F) cortex. Solid bars indicate subsystems with C removed. Striped bars indicate Φ* for subsystems where C data are replaced with gaussian noise. (**B**) Φ* (± SE) calculated for system containing multiple electrode contacts in C (striped) or only a single contact in C (solid) during wakefulness, sleep and anesthesia (Anes).

**Fig. S4.**
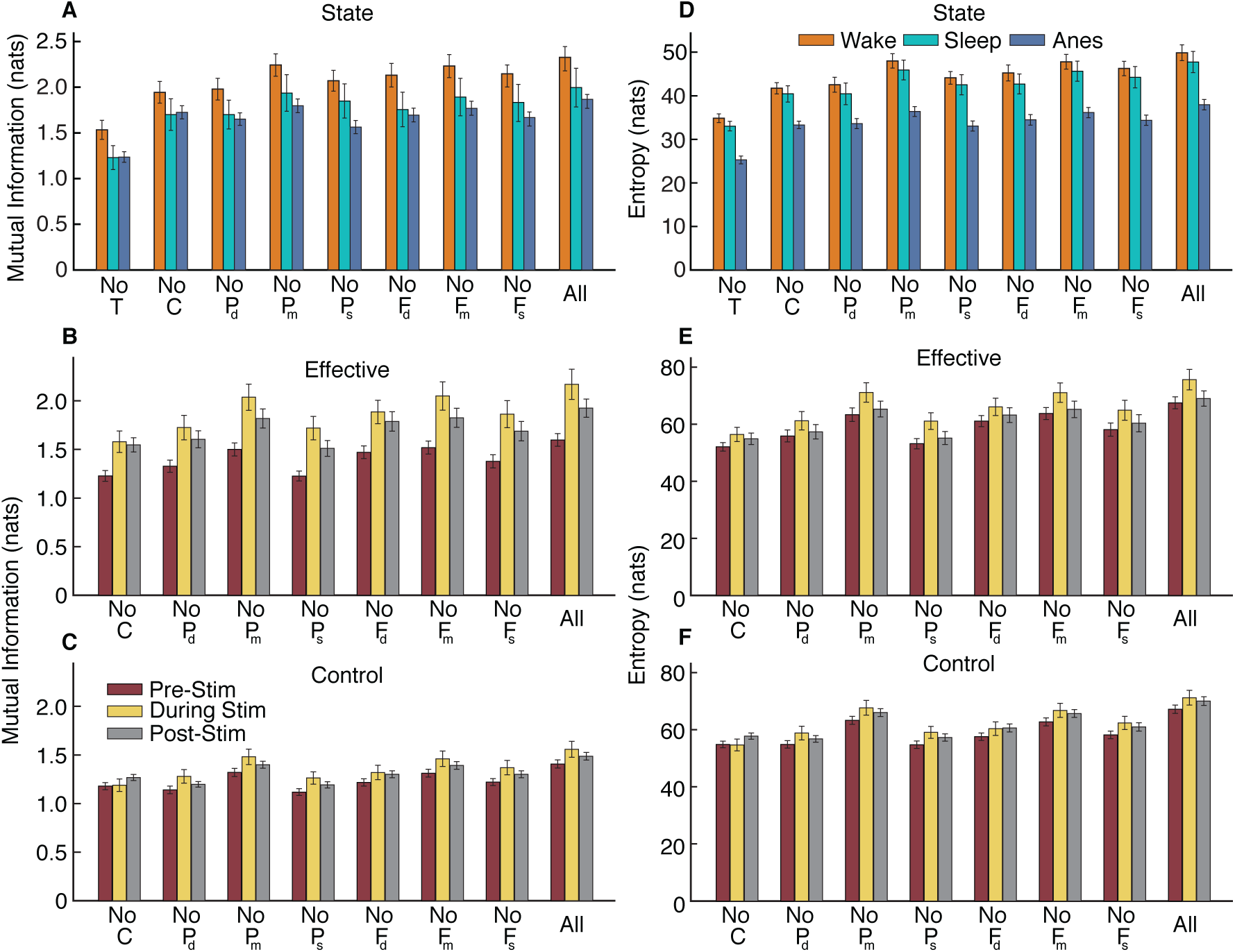
Entropy and mutual information vary with conscious state, but less so with thalamic stimulations that change consciousness level. (**A**) Average mutual information (± SE) for different subsystems absent thalamus (T), caudate nucleus (C), superficial (s), middle (m) or deep (d) layers of parietal (P) or frontal (F) cortex, or for the full system of 8 parts (All; CTP_d_P_m_P_s_F_d_F_m_F_s_), during wakefulness (orange), sleep (teal) and anesthesia (blue). (**B** and **C**) Average mutual information (± SE) prior to (burgundy), during (yellow) or after (gray) effective (**B**) or control (**C**) thalamic stimulation. (**D**) Average entropy (± SE) for different subsystems and full system, during wakefulness (orange), sleep (teal) and anesthesia (blue). (**E** and **F**) Average entropy (± SE) prior to (burgundy), during (yellow) or after (gray) effective (**E**) or control (**F**) thalamic stimulation.

**Fig. S5.**
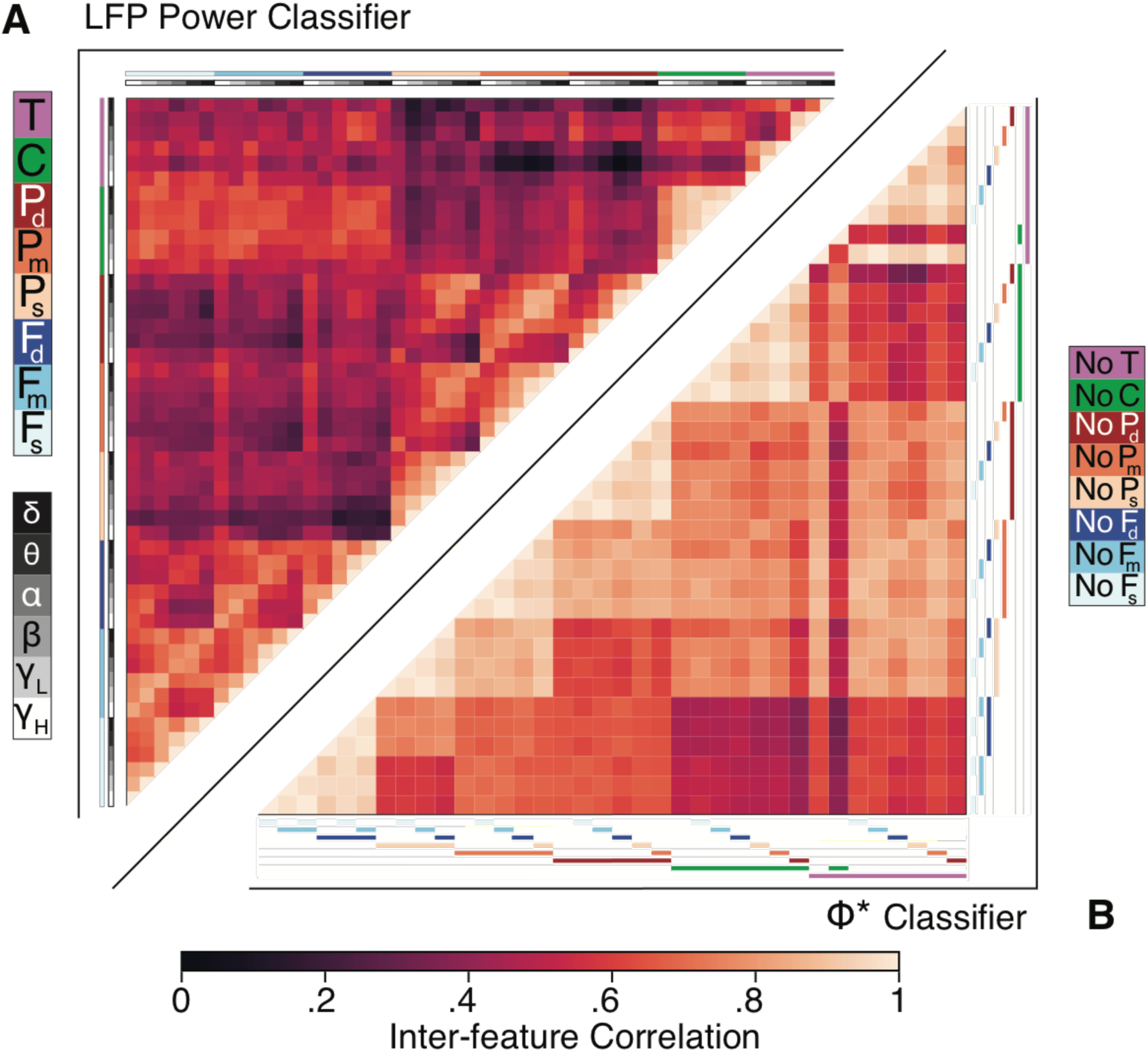
Limited inter-feature correlation for both the LFP power and Φ* classifiers justifies MDA analyses of feature importance. (**A**, top left) Inter-feature correlation for the LFP power classifier. Features are LFP power at delta (δ = 0-4 Hz), theta (θ = 4-8 Hz), alpha (α = 8-15 Hz), beta (β = 15-30 Hz), low gamma (γ_L_ = 30-60 Hz) and high gamma (γ_H_ = 60-90 Hz) within thalamus (T), caudate (C), superficial (s), middle (m) or deep (d) layers of parietal (P) or frontal (F) cortex. (**B**, bottom right) Inter-feature correlation for the Φ* classifier. Features are subsystems absent thalamus (T), caudate (C), superficial (s), middle (m) or deep (d) layers of parietal (P) or frontal (F) cortex.

**Fig. S6.**
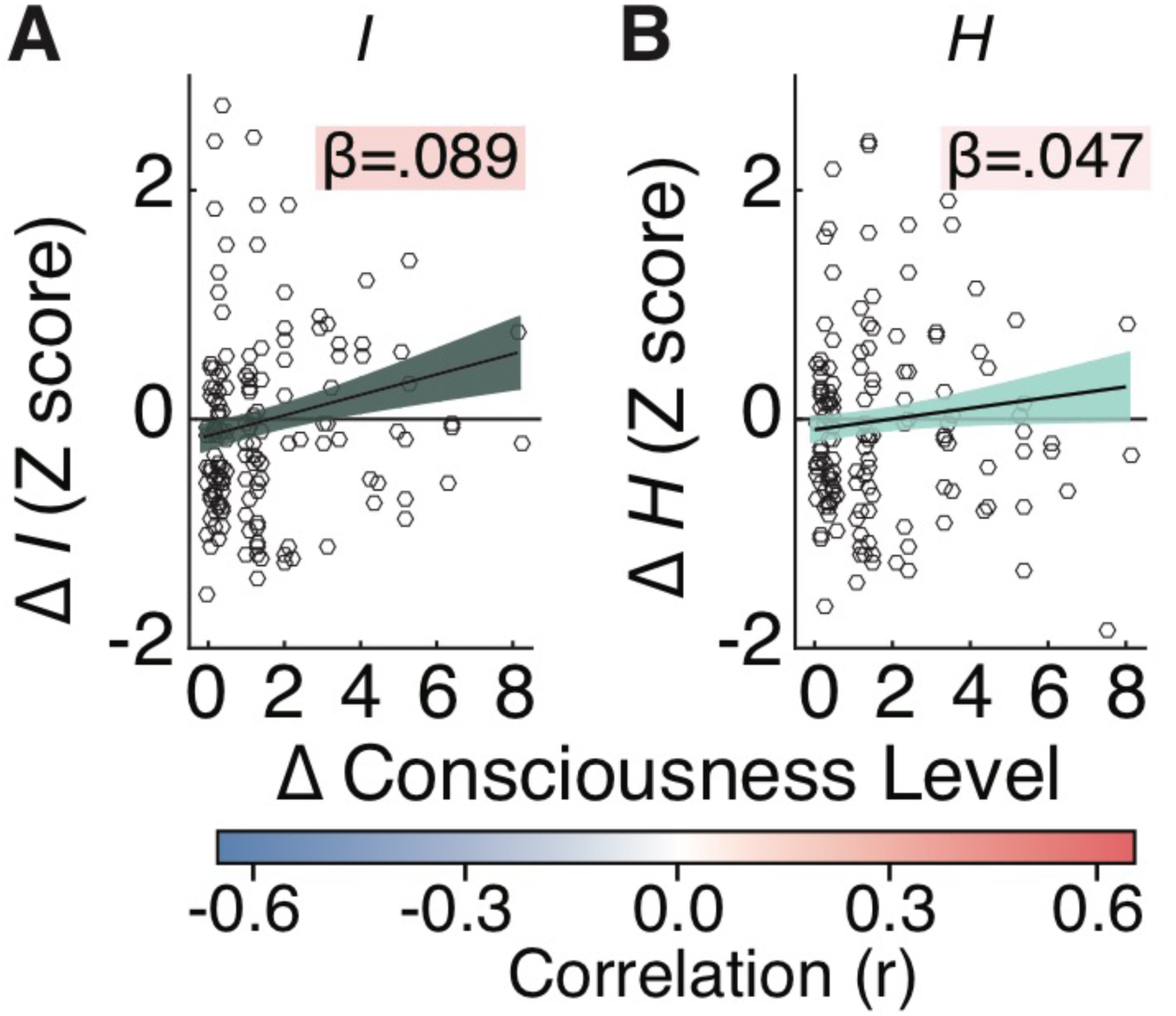
Normalized entropy (*H*) and mutual information (*I*) do not correlate well with stimulation-induced changes in level of consciousness. (**A**) Regression estimate for the effect of stimulation-induced changes in level of consciousness (during stim – pre) on normalized *I* (Z score, during stim – pre). (**B**) Regression estimate for the effect of stimulation-induced changes in level of consciousness (during stim – pre) on normalized *H* (Z score, during stim – pre). Points show values for each stimulation event. Lines show regression fit (± SE) with reported slope (β). Color behind β indicates r value.

**Fig. S7.**
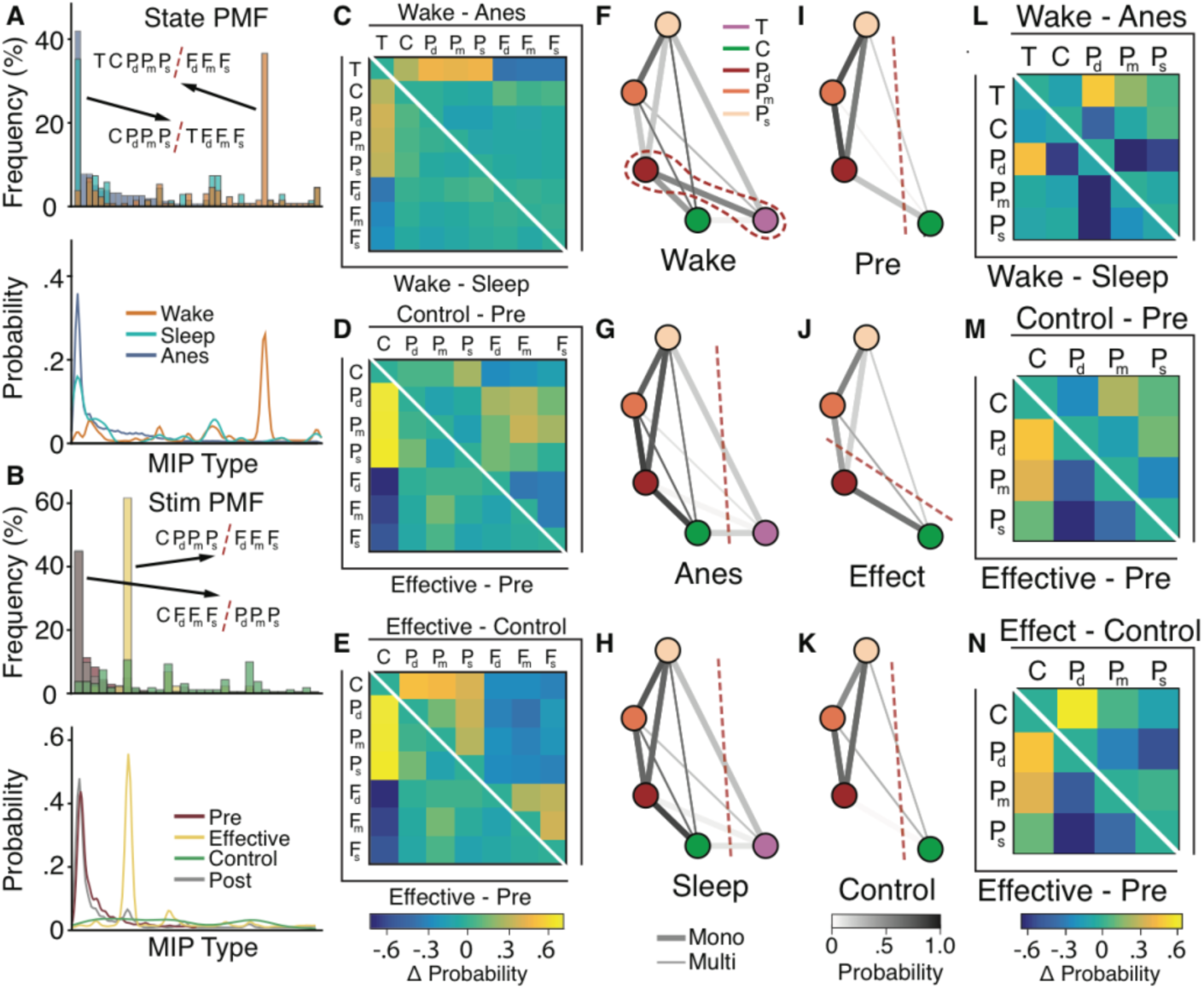
Minimum information partitions (MIPs) show that consciousness coincides with increased affiliation of parietal deep layers and subcortical areas. (**A**) Probability mass functions (PMF) of MIP types for each state, normalized by state, and aligned to the anesthesia condition, with accompanying kernel density estimates (**B**) PMF of MIP types for each stimulation condition, normalized by condition, aligned to pre-stimulation, with the accompanying kernel density estimates. Identified MIPs represent the peak of the distribution. Red dashed lines indicate the “cut” separating partitioned areas (thalamus = T, caudate = C, superficial, middle, deep parietal = (P_s_, P_m_, P_d_), and frontal = (F_s_, F_m_, F_d_),). (**C-E**) Differences in pairwise probability of brain areas associating on the same side of the MIP (from Fig. 4, F-K) for state and stimulation comparisons relevant for consciousness: (**C**) wake – sleep compared to wake – anesthesia; (**D**) effective stimulation – pre-stimulation compared to control stimulation – pre-stimulation; and (**E**) compared to effective stimulation – control stimulation. Color scale shows change in probable affiliation of brain areas in same MIP side. (**F-K**) Pairwise probability (gray scale lines) of brain areas associating on the same side of the MIP for the 5-part system (TCP_s_P_m_P_d_) during (**F**) wakefulness, (**G**) anesthesia and (**H**) sleep, and for the 4-part system (TCP_s_P_m_P_d_) (**I**) prior to, (**J**) during effective, and (**K**) during control stimulation. Gray scales with probability of association. Thick lines show monosynaptic (Mono) anatomical paths (in at least one direction); thin lines show multisynaptic (Multi) paths. Dashed red lines show dominant MIP when applicable. (**L-N**) Differences in pairwise probability of brain areas associating on the same side of the MIP (from F-K) for state and stimulation comparisons relevant for consciousness: (**L**) Wake – sleep compared to Wake – anesthesia; (**M**) effective stimulation – pre-stimulation compared to control stimulation – pre-stimulation; and (**N**) compared to effective stimulation – control stimulation. Color scale shows change in probable affiliation of brain areas in same MIP side.

**Table S1.**
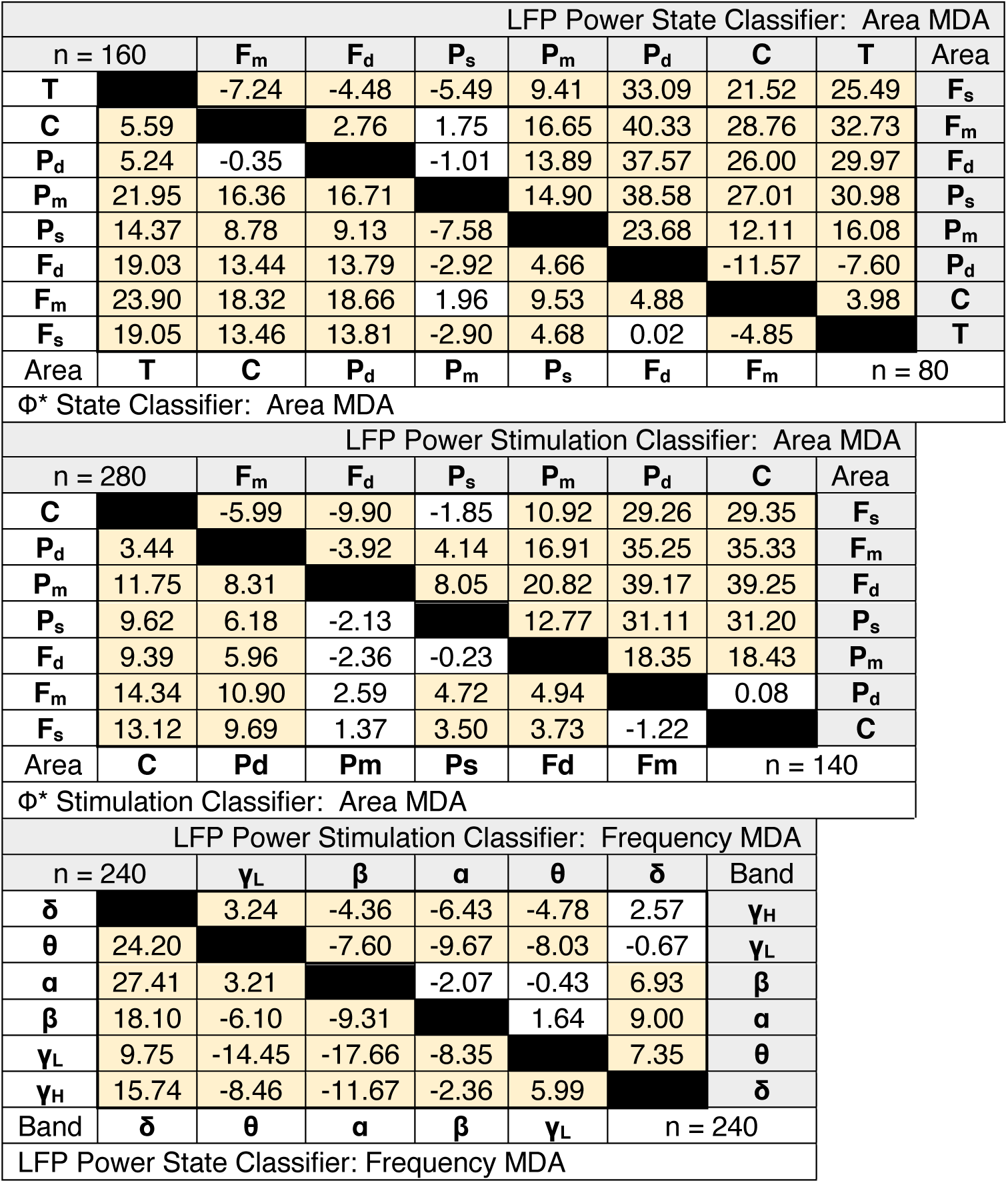
Statistical results for all mean decrease in accuracy (MDA) analyses (related to Fig. 1, F-H and Fig. 2, H-J). Values are t statistics for pairwise comparisons of MDA related to features within all classifiers. Positive values reflect the column feature higher than the row feature on the same side of the diagonal with the same color. Yellow shading indicates statistical significance (p < 0.05) after controlling for multiple comparisons. Classifiers are separated along black diagonals according to state or stimulation comparisons, or measure (LFP power or Φ*) comparisons. For example, for the LFP Power stimulation classifier (middle gray), P_s_ (column feature) leads to a significantly higher MDA than F_m_ (row feature, t = 4.14), but an insignificantly lower MDA than F_s_ (t = −1.85).

**Table S2.** Statistical results for all LFP power linear regression correlation analyses (related to Fig. 3, B-H). Values are β slopes and t statistics for the regression of changes in normalized power of each brain area (during – pre-stimulation) and change in level of consciousness (during – pre-stimulation). Columns, frequency bands; rows, brain areas: caudate (C), superficial (s), middle (m) or deep (d) layers of parietal (P) or frontal (F) ROIs. Positive β slopes or t values indicate an increase in power related to an increase in consciousness level. No tests were statistically significant (p > .05).

| Linear Regression ($\Delta$ Power ∼ $\Delta$ consciousness level) | | | | | | | | | | | | |
| --- | --- | --- | --- | --- | --- | --- | --- | --- | --- | --- | --- | --- |
| n = 163 | $\delta$ | | $\theta$ | | $\alpha$ | | $\beta$ | | $\gamma_L$ | | $\gamma_H$ | |
| | $\beta$ | t | $\beta$ | t | $\beta$ | t | $\beta$ | t | $\beta$ | t | $\beta$ | t |
| **C** | -0.05 | -1.30 | -0.05 | -1.48 | -0.05 | -1.45 | -0.02 | -0.57 | 0.00 | 0.00 | -0.04 | 1.00 |
| **P<sub>d</sub>** | 0.07 | 1.56 | 0.04 | 0.95 | 0.02 | 0.52 | 0.02 | 0.42 | 0.02 | 0.50 | 0.01 | 0.33 |
| **P<sub>m</sub>** | 0.02 | 0.33 | 0.02 | 0.32 | 0.01 | 0.30 | 0.00 | 0.12 | 0.02 | 0.43 | 0.00 | -0.02 |
| **P<sub>s</sub>** | -0.05 | -0.98 | -0.05 | -1.07 | -0.06 | -1.36 | -0.05 | -1.36 | -0.01 | -0.25 | -0.03 | -0.68 |
| **F<sub>d</sub>** | 0.05 | 1.08 | 0.04 | 0.88 | 0.03 | 0.67 | 0.04 | 0.85 | 0.03 | 0.92 | 0.04 | 1.10 |
| **F<sub>m</sub>** | -0.01 | -0.31 | 0.00 | -0.11 | -0.01 | -0.14 | 0.01 | 0.15 | 0.03 | 0.84 | 0.01 | 0.30 |
| **F<sub>s</sub>** | 0.06 | 1.27 | 0.05 | 1.09 | 0.03 | 0.62 | 0.02 | 0.46 | 0.02 | 0.48 | 0.03 | 0.72 |

**Table S3.** Statistical results for pairwise comparisons of the influence of brain area subsystem membership on Φ* (related to Fig. 4, A and C). Values are t statistics for pairwise comparisons of Φ* generated by subsystems containing the column area but not the row area and vice versa. Positive values when the column area-containing subsystems had higher Φ* than the row area-containing subsystems on the same side of the diagonal with the same color. Yellow shading indicates statistical significance (p <.05) after controlling for multiple comparisons. State comparisons correspond to areas on white background, whereas stimulations correspond to areas on gray background. For example, for the Φ* stimulation data (gray), C (column area) yields significantly higher Φ* than P_s_ (row area, t = 5.55), but insignificantly lower Φ* than P_d_ (row area, t = −2.32).

| Φ* Stimulation: System Membership |  |  |  |  |  |  |  |  |
| --- | --- | --- | --- | --- | --- | --- | --- | --- |
| n = 372 |  | F<sub>m</sub> | F<sub>d</sub> | P<sub>s</sub> | P<sub>m</sub> | P<sub>d</sub> | C | Area |
| T |  | 0.16 | 2.92 | 6.95 | 5.26 | 15.10 | 13.18 | F<sub>s</sub> |
| C | 14.33 |  | 2.74 | 6.77 | 5.08 | 14.95 | 13.11 | F<sub>m</sub> |
| P<sub>d</sub> | 16.14 | 1.29 |  | 3.97 | 2.29 | 11.95 | 10.49 | F<sub>d</sub> |
| P<sub>m</sub> | 20.72 | 7.14 | 5.90 |  | -1.62 | 7.36 | 5.55 | P<sub>s</sub> |
| P<sub>s</sub> | 21.58 | 7.69 | 6.47 | 0.51 |  | 8.95 | 7.29 | P<sub>m</sub> |
| F<sub>d</sub> | 21.79 | 8.29 | 7.11 | 1.27 | 0.76 |  | -2.32 | P<sub>d</sub> |
| F<sub>m</sub> | 23.54 | 9.47 | 8.23 | 2.30 | 1.79 | 1.02 |  | C |
| F<sub>s</sub> | 23.54 | 9.29 | 8.05 | 2.10 | 1.59 | 0.82 | -0.20 | n = 378 |
| Area | T | C | P<sub>d</sub> | P<sub>m</sub> | P<sub>s</sub> | F<sub>d</sub> | F<sub>m</sub> |  |
| Φ* State: System Membership |  |  |  |  |  |  |  |  |

